# Hair Bundle Degeneration is a Key Contributor to Age-Related Vestibular Dysfunction

**DOI:** 10.1101/2025.02.02.636113

**Authors:** Samadhi Kulasooriya, Huizhan Liu, Sarath Vijayakumar, Celia Bloom, Shu Tu, Benjamin J. Borgmeier, Mi Zhou, Litao Tao, Bechara Kachar, David Z. He

**Author notes:** **Corresponding authors:** David Z. He, Samadhi Kulasooriya. **Teaser:** Hair bundle degeneration and age-related vestibular dysfunction.

## Abstract

Age-related vestibular dysfunction (ARVD) is a prevalent, debilitating condition in the elderly. The etiology and molecular mechanisms are poorly understood. We focused on mechanosensitive hair cells (HCs) as they are vulnerable to aging. Using single-cell RNA-seq transcriptomes of young and old mouse vestibular HCs, we show that aging vestibular HCs display both universal molecular signatures, such as genomic instability, mitochondrial dysfunction, and impaired proteostasis, and cell type-specific changes associated with deterioration of hair bundles and mechanotransduction. In alignment with transcriptomic findings, imaging and electrophysiological recordings from aged vestibular sensory epithelia confirmed the degeneration of the hair bundles and a reduction in mechanotransduction. Importantly, this deterioration of hair bundles and vestibular function precedes HC loss, highlighting impaired mechanotransduction as a key contributor to ARVD. Furthermore, molecular and cellular changes associated with aging signatures are less pronounced in vestibular HCs than in cochlear HCs, underscoring tissue-specific age-related differences between the two sensory epithelia in the inner ear.

## Introduction

Aging research has experienced unprecedented advances in recent years, revealing that genetic and molecular processes conserved through evolution may partially govern the pace of aging (*1*). Aging is the progressive deterioration of physiological integrity, leading to impaired function and enhanced vulnerability to disease and death. The molecular processes of aging are characterized by the universal hallmarks of aging, including increased cellular senescence, genomic instability, epigenetic alterations, mitochondrial dysfunction, loss of proteostasis, and inflammaging (*1*, *2*). The interconnected and interdependent nature of these processes leads to slow age-related structural changes and functional decline. Similar to other sensory organs, biological aging negatively impacts the inner ear, resulting in age-related hearing loss (ARHL) and vestibular dysfunction (ARVD). While numerous studies have focused on ARHL, ARVD remains relatively understudied, partially due to its multifaceted etiology (*3–8*). ARVD is characterized by the gradual loss of bilateral vestibular function accompanied by interruptions to visual and proprioceptive inputs. The damaging impact of age-related vestibular sensory decline manifests itself in an exponential increase in geriatric dizziness and injurious/fatal falls (*9*). According to the National Center for Health Statistics and National Institutes of Health, the prevalence of ARVD in the US is 75.3%. It is the 6^th^ leading cause of death and accounts for 50% of all accidental deaths among the elderly population (*5*, *10*).

While the central neural pathways in the vestibular system play a role in ARVD, age-related degeneration of vestibular hair cells (HCs) and neurons in the vestibular periphery contributes to ARVD (*11*). The vestibular system plays a crucial role in sensing the direction, acceleration, and spatial orientation of the head in response to gravity to maintain the upright posture and balance of the body. It consists of three crista ampullaris that detect the angular motions and two otolith organs (utricle and saccule) that detect linear motions of the head (*12*). Vestibular sensory epithelia consist of two types of mechanosensitive HCs, type I and type II. HCs convert mechanical stimuli into electrical signals. Mechanotransduction, the first step in the vestibular signal processing, is mediated by the mechanotransduction apparatus in the hair bundle comprising tightly linked stereocilia and a kinocilium (*13*, *14*). Therefore, damage or degeneration of hair bundles will lead to vestibular functional decline (*11*, *13–16*). Interestingly, studies from human and murine models indicate an earlier and more pronounced auditory functional and morphological decline compared to the vestibular system, highlighting tissue-specific differences between these sensory organs (*7*, *9*). However, molecular mechanisms driving these divergent aging processes remain unknown. This fundamental lack of understanding of the molecular basis of vestibular aging remains a significant barrier to developing targeted therapeutic interventions to mitigate ARVD (*17*).

While previous studies have examined age-related physiological and morphological changes in the vestibular system, no studies have been performed to assess age-related changes of vestibular HCs at the molecular level. Therefore, it is still unclear whether vestibular HCs follow the universal blueprint of aging as seen in neurons and cochlear HCs, and whether vestibular HCs also exhibit cell-type-specific aging signatures. In the current study, we comprehensively examined the aging of the vestibular system across functional, morphological, and molecular levels. Transcriptomic studies have been at the forefront of revealing the molecular underpinnings of development, aging, and disease conditions in model organisms and humans. Utilizing single-cell RNA sequencing (scRNA-seq), we characterized universal and cell-type-specific aging signatures in vestibular HCs from young and aged mice. The cell type-specific aging signatures are primarily associated with the degeneration of hair bundles and the mechanotransduction apparatus. Concordant with that, electrophysiology and imaging revealed that vestibular functional decline and bundle degeneration precede HC loss, highlighting the degeneration of the hair bundles as a key contributor to ARVD. The comparison between vestibular and cochlear HCs shows that the age-related molecular and cellular changes are less pronounced in vestibular HCs than in cochlear HCs, underscoring the subtle differences in aging between the two sensory epithelia in the mammalian inner ear. Our data also provide a foundational transcriptomic resource to advance the understanding of inner ear aging and the development of targeted therapeutic strategies.

## Results

### Age-related functional and morphological changes in the vestibular HCs

Prior studies have reported various age-related functional and structural changes in the vestibular end organs (*5*, *6*, *18*, *19*). In this study, we utilized CBA/J mice, which exhibit progressive age-related changes in the inner ear in the later stage of life, similar to humans (*20*, *21*). To examine the age-related functional decline in the vestibular system, we measured vestibular sensory evoked potentials (VsEP) from 2-2.5 months (young) and 22-24 months (old) mouse cohorts. This test measures electrical signals generated by the vestibular sensory epithelia in response to rapid linear head accelerations by evaluating sensitivity (threshold), strength of the response (amplitude), and time taken by the signal to reach the recording electrode after stimulus initiation (latency) (*22*, *23*). Consistent with previous findings, our results indicated an age-related decline in the vestibular function, as evidenced by a significant decrease in the response amplitude and an increase in the threshold and latency (Fig.1, A to C). We also measured endolymphatic potential (ELP), which is essential for maintaining the ionic gradient required for mechanotransduction (*24*), and it typically ranges from +1 to +11 mV (*25–27*). Our findings revealed no age-related change in ELP, indicating maintenance of a stable electrochemical environment (Fig.1D). Next, we examined the age-related morphological alterations in vestibular HCs. Previous studies have reported an age-related loss of vestibular HCs in both mice and humans (*6*, *16*, *28*, *29*), with more loss of type I HCs than type II HCs (*6*), while some studies indicate no HC loss (*30*). Thus, to re-examine the age-related HC loss, we immunolabeled young and old utricles with HC-specific marker MYO7A and type II HC-specific marker SOX2 (*31–33*). Our findings showed no age-related changes in type I and II HC density in the striolar and extrastriolar regions (Fig.1, E and F, fig. S1). Cellular hypertrophy and atrophy are commonly seen in aging cells, including cochlear HCs and neurons (*33*, *34*). To further assess the age-related cytological changes in vestibular HCs, we performed a histological analysis (Fig.1, G to I). While we observed cellular hypertrophy and atrophy in some HCs, there were no significant changes in cell length (Fig.1, H and I).

**Fig. 1.**
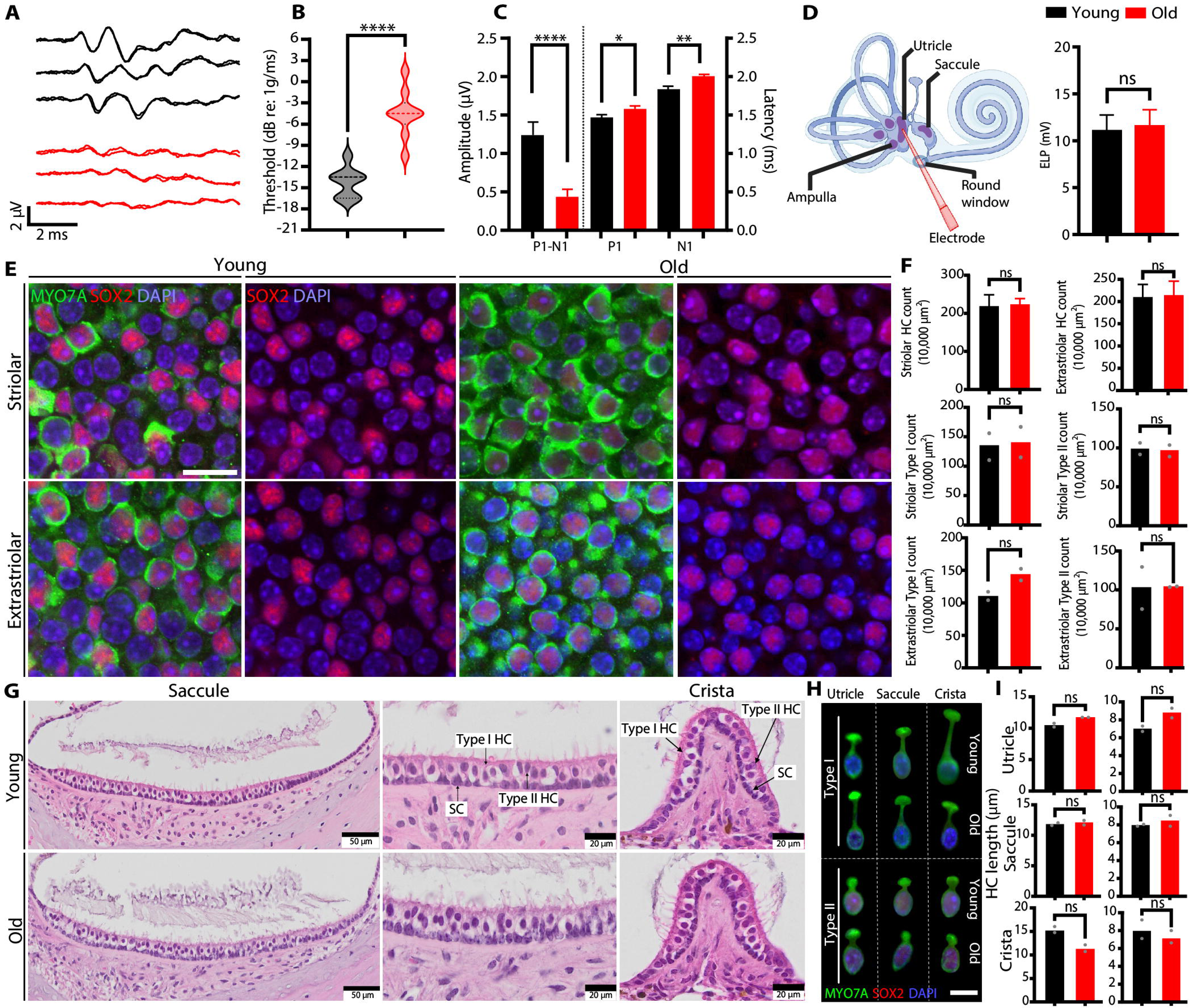
Age-related functional and cellular changes in the vestibular HCs. (**A** to **C).** Measurements of VsEP from the young and old mice (n = 8 per age group). (A) 3 representative wave forms from each age group depicting the reduction in response amplitude, (B) elevation of threshold in the old animals compared to young animals, and (C) quantification of reduction in response amplitude (left) and increase in latency in the old group compared to the young group (right). P1 and N1 indicate the first positive and negative peaks. **(D)** Graphical illustration of measurement of ELP using a glass capillary microelectrode (D), and ELP magnitude quantifications (n=6 per age group). **(E)** Representative images of young and old striolar and extrastriolar sensory epithelia (cropped region) from utricle whole mounts immunolabeled with anti-MYO7A and anti-SOX2, Nuclei were stained with DAPI. Scale bar, 10 µm. **(F)** Quantifications of total HC density (n=4), as well as type I and II HC density in striolar and extrastriolar regions (n=2). **(G)** Representative images of young and old H&E sections indicating type I and II HCs from the saccule and cristae. **(H)** Confocal virtual sectioning images of some individual HCs demonstrating cellular hypertrophy (elongation) and atrophy (shortening) in the three vestibular end organs acquired by the ortho slicer tool in Imaris application. Scale bar, 10 µm. **(I)** Quantification of HC lengths in the three vestibular end organs from histological analysis. Individual cell measurements of type I and II HCs were taken along the sensory epithelia (utricle, saccule, and cristae) from two technical replicates per biological replicate (n=2 per age group). Data are shown as mean ± SEM, *p <0.05, **p <0.01, ****p<0.0001, ns - non-significant by unpaired t-test.

We utilized high-resolution confocal imaging to examine the age-related changes in the hair bundle. Stereocilia were labeled with phalloidin, and kinocilia were immunostained with anti-acetylated-β-tubulin. We observed signs of stereociliopathy, including complete and partial loss of stereocilia leading to bundle thinning and shortening, and fusion of stereocilia, alongside degeneration of kinocilia (Fig. 2A). While cochlear HCs lose the kinocilia during maturation, vestibular kinocilia are maintained in adult HCs and play a crucial role in hair bundle polarity, determining directional sensitivity and connecting the hair bundle to the overlaying otolithic membrane (*35*, *36*). We observed signs of kinociliary degeneration, including kinocilia fusion (fuses with neighboring kinocilia or stereocilia), and shortening, tip or mid-shaft bulb formation, aberrant looping, and knotting (Fig. 2A). These degenerative changes increased with aging in both the utricle and cristae (Fig. 2B). We also utilized scanning electron microscopy (SEM) to examine the ultrastructure of the hair bundle, and we observed various signs of bundle degeneration (Fig. 2C). Collectively, these findings indicate various age-related changes, particularly in the hair bundle, that may impair mechanotransduction and sensory perception, leading to vestibular functional decline as indicated by VsEP.

**Fig. 2.**
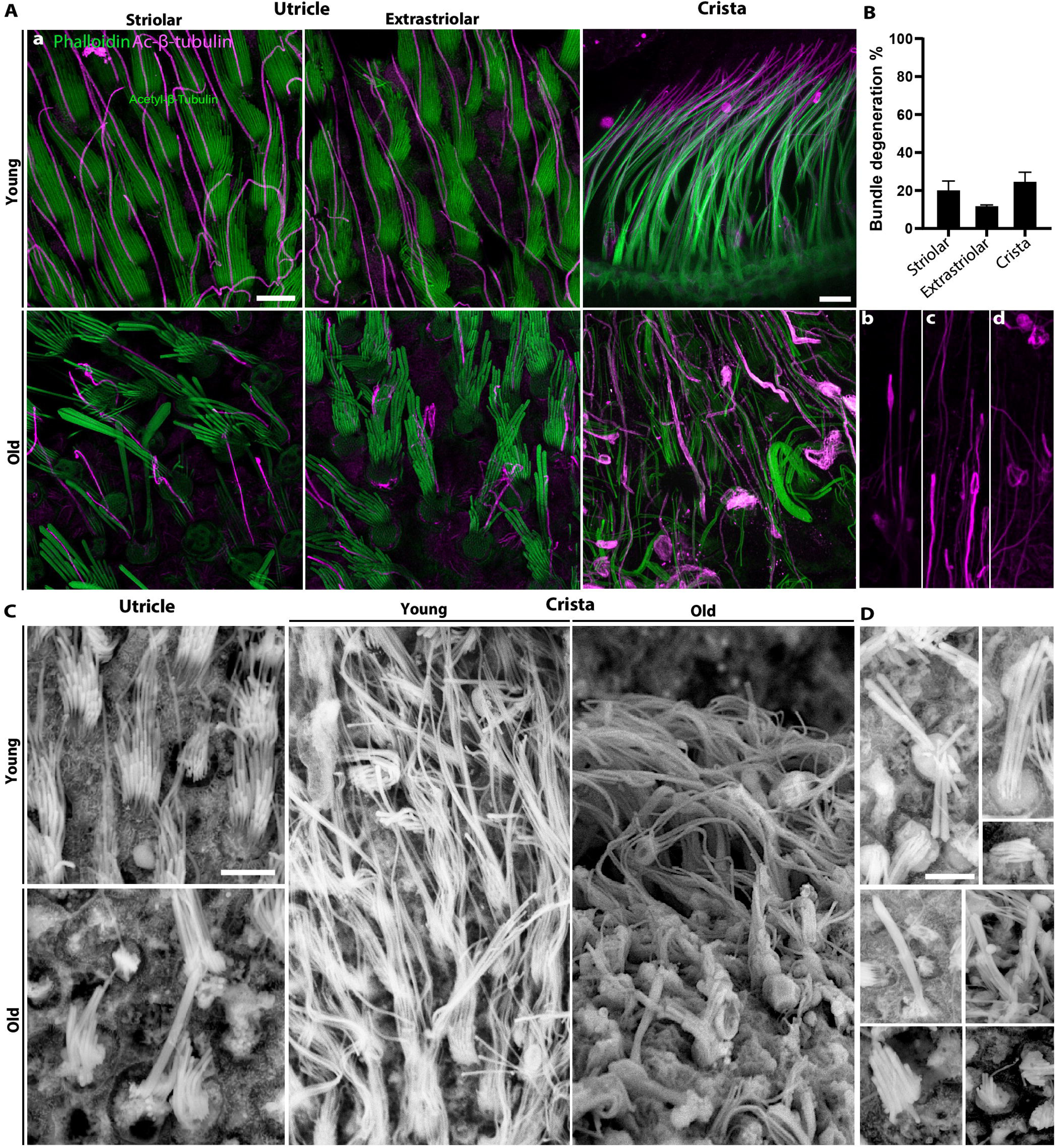
Age-related changes in the hair bundle. **(A)** High-resolution confocal microscopic images of young and old utricle and cristae, indicating stereocilia and kinocilia degeneration with aging (a). Stereocilia were stained with Phalloidin, and kinocilia were immunolabeled with anti-acetylated-β-tubulin. Scale bar, 5 µm. Lower right representative images indicate the kinociliary bulb formation (b), fusion (c), looping, and knotting (d) with aging. **(B)** Quantifications of degenerative changes in young and old utricle and cristae (n=4). The total number of bundles and bundles with degenerative signs were quantified from two different areas in striolar and extrastriolar regions per biological replicate in the two age groups and presented as a percentage. Data shown as mean ± SEM. **(C)** Representative SEM images of hair bundles from young and old utricle and cristae indicating signs of degeneration. Scale bar, 5 µm. **(D)** Examples of various signs of bundle degeneration. Scale bar, 5 µm. All images were taken from regions with obvious signs of bundle degeneration.

### Cellular diversity in young and old vestibular end organs

The molecular mechanisms underlying age-related vestibular degeneration are poorly understood (*37*). To unveil this, we conducted single-cell RNA-sequencing (scRNA-seq) of inner ears harvested from 10-week- and 22-month-old CBA/J mice. The sequencing reads were aligned to the mouse reference genome (mm10), followed by standard quality control metrics and downstream processing steps, including dimensional reduction and unsupervised clustering (Fig.3A, fig. S2) (*38*). Cell clusters were annotated based on known marker gene expression and gene signatures (Fig.3, B and C). We examined the cellular diversity in the vestibular sensory epithelia of young and old samples, encompassing type I and II HCs, supporting cells, and blood, neuronal, immune, and epithelial cell populations. HCs were identified based on the expression of markers such as *Pou4f3, Myo7a, Slc17a8,* and *Otof* (*39*, *40*), and type I and II HC types were differentiated using *Sox2* (*31*, *32*) (Fig.3D, E, G, H). Based on these markers, 698 type I and 129 type II HCs from young samples and 1651 type I and 230 type II HCs from old samples were isolated for the downstream analysis (Fig.3, F and I).

**Fig. 3:**
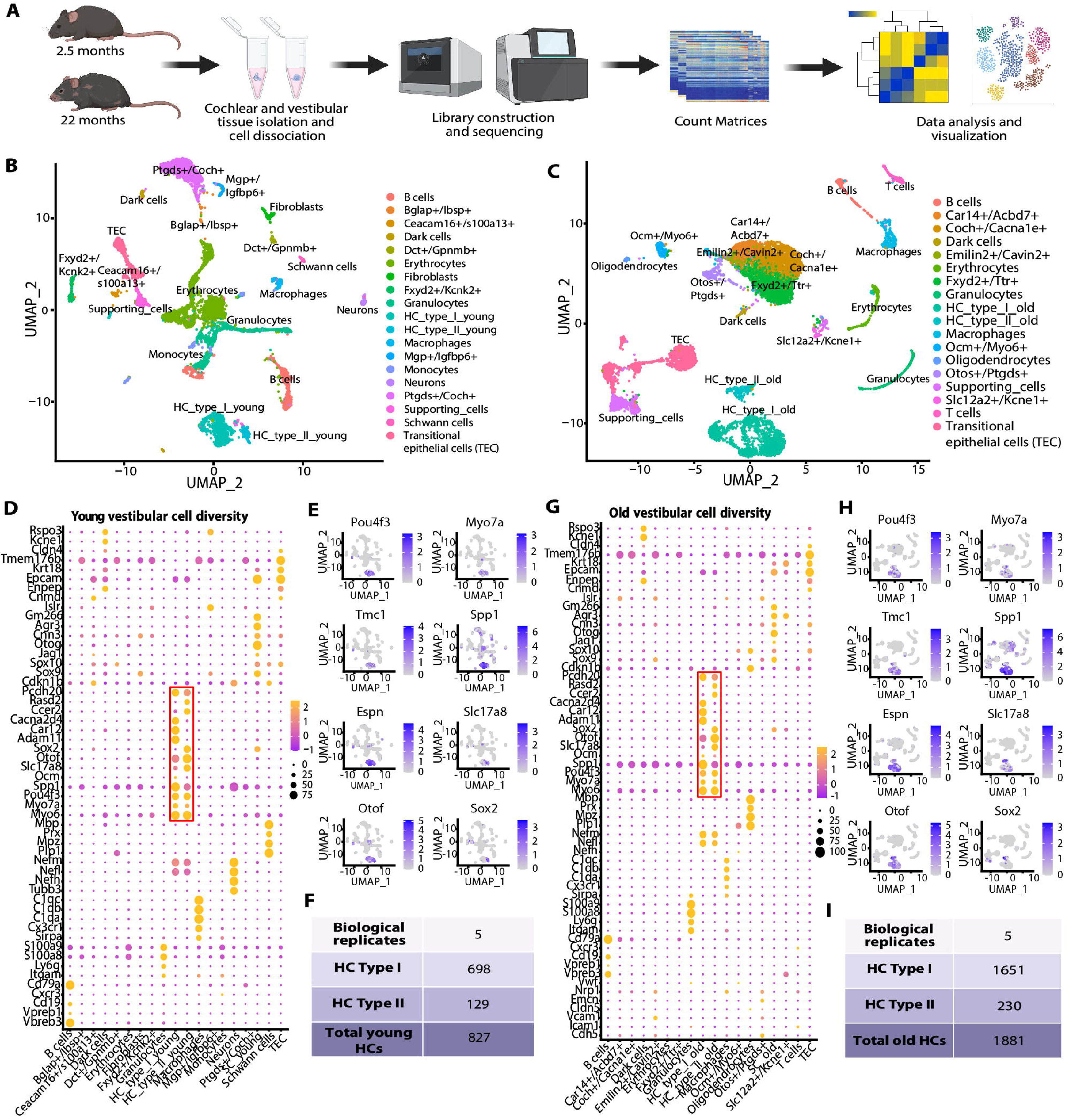
Vestibular cell type distribution and HC cluster annotation. **(A)** An illustration of the experimental workflow of scRNA-sequencing, data analysis, and visualization. **(B and C)** Uniform manifold approximation and projection (UMAP) plot showing the distribution of different cell types in young and old vestibular samples, respectively. This dimensional reduction method uses k-nearest neighbors (KNN) and the Euclidean distance algorithm to construct a low-dimensional graph for visualization. **(D and G)** Dot plots indicating the expression of known marker genes and gene signatures used for cluster annotation. The dot size represents the percentage of cells expressing the genes from each cluster, whereas the color indicates the expression level. Expression levels are normalized via z-score normalization. Thus, the average expression is zero, and positive or negative values indicate expression above or below average. **(E and H)** Feature plots demonstrating the HC-specific pan marker gene expression in the HC clusters identified in the UMAPs. **(F and I)** Number of type I and II HCs identified from young and old vestibular samples for the downstream analysis (n=7-8 mice per biological replicate).

### Universal hallmarks of aging

Cellular aging typically follows universal aging programs (*1*, *2*). Prior studies have demonstrated that cochlear HC aging is associated with universal hallmarks such as mitochondrial dysfunction, oxidative stress, apoptosis, and genomic instability (*17*). Thus, we speculated that aging vestibular HCs may also exhibit universal hallmarks. We aggregated single-cell raw counts to generate pseudobulk expression matrices of young and old vestibular HCs (table S2), and the gene expression levels related to universal hallmarks were analyzed using genes identified from multiple resources as described in the methods (Fig.4A). Cellular senescence is characterized by the permanent cessation of the cell cycle. Senescent cells adopt the characteristic senescence-associated secretory phenotypes (SASP), which exert deleterious effects on cellular function over time and are recognized as a hallmark of cellular aging (*1*, *2*, *41*). Our data indicated a decrease in cell cycle regulators (*Cdk2, Cdk4, Cdk6*), followed by an increase in cyclin-dependent kinase inhibitors (*Cdkn1a, Cdkn1b*), and degradation of cyclins (*Fzr1*), indicating cell cycle arrest. Moreover, we noted an increase in senescence induction (*Ypel3*), a decrease in laminins (*Lmna, Lmnb1*), and enrichment of various SASPs that are involved in inflammation (*Cxcl14, Nfkb2, B2m*), senescence-associated heterochromatic foci (*H3f3a, Hira, Ubn1*), hypoxia, and angiogenesis (*Vegfa, Hif1a, Nos3*), indicating increased senescence in old HCs relative to young HCs. The imbalance between the production and elimination of reactive oxygen species (ROS) results in oxidative stress (*1*, *2*). We observed an increased expression of genes involved in mitochondrial respiration, the major contributor of ROS generation (*Cox4i1, Cox6a1, mt-Co1, mt-Cytb, Ndufs2*), antioxidant response and detoxification (*Mt1, Sod1, Prdx1, Gpx4, Keap1, Nrf1*), mitochondrial fission and fusion (*Fis1, Mfn1, Mfn2*), stress response (*Bcl2, Atf4, Cflar, Hif1a, Atf4, Atf5*) and mitochondrial quality control (*Lonp1, Trap1*), suggesting age-related increase in oxidative stress followed by a compensatory increase in antioxidant mechanisms. In addition, we noticed a decrease in key genes involved in the detoxification of hydrogen peroxide, such as *Gpx1, Prdx1, Sod3*, and *Cat*, indicating partial ROS clearance.

**Fig. 4.**
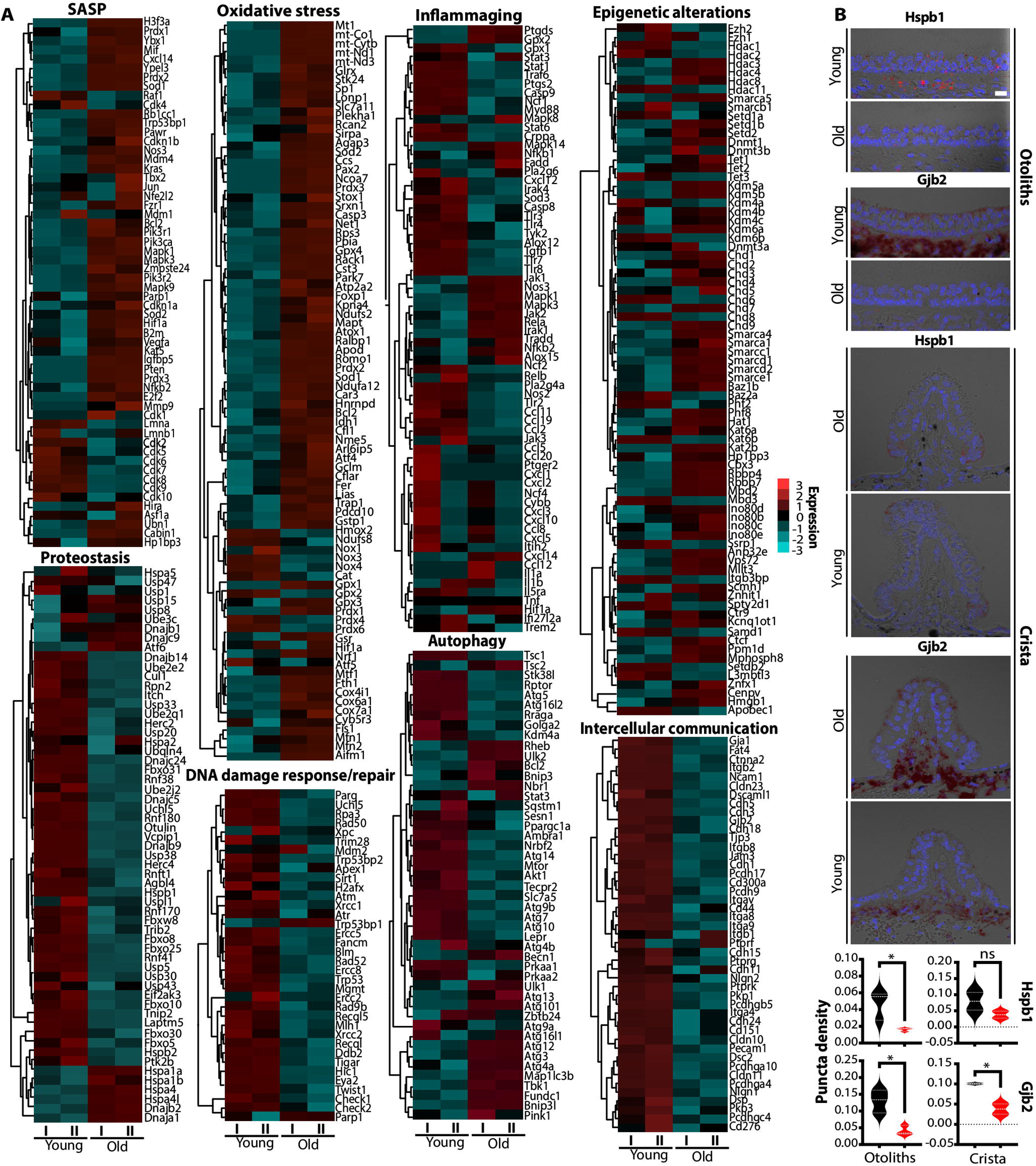
Universal hallmarks of aging in vestibular HCs. **(A)** Complex heatmaps indicating the enrichment of genes associated with universal hallmarks of aging, including senescence and SASP, loss of proteostasis, oxidative stress, autophagy, inflammaging, DNA damage repair, intercellular communication, and epigenetic alterations in the old vestibular HCs compared to young HCs. Single-cell counts were aggregated. Pseudo-bulk gene expression values were then scaled and log2-transformed. Expression values were centered by Cluster 3.0, and heatmaps were generated using JAVA TreeView. The color scale indicates the expression level. **(B)** RNAscope *in situ* hybridization of *Hspb1* (proteostasis) and *Gjb2* (intercellular communication) in young and old otoliths (n=3) and crista (n=2). Scale bar, 10 µm. The puncta density for each gene was quantified using FIJI. Data shown as mean ± SEM, *p <0.05 by unpaired t-test.

Age-related gradual increase in chronic low-grade inflammation is termed inflammaging (*1*, *2*). We noticed a steady increase in pro-inflammatory genes (*Ptgds, Cxcl14, Ccl12, Il1a, Nfkb1, Jak1, Stat3, Tradd, Fadd, Irak1, Mapk1, Mapk14*) and downregulation of some genes (*Tgfb1, Cxcl12, Tlr3, Tlr4, Myd88, Irak4, Ccl2, Ccl5, Ccl11*), indicating potential maintenance of a prolonged low-grade inflammatory state followed by reduced immune surveillance and response. Epigenetic dysregulation increases with aging (*42*). Our data demonstrated age-related epigenetic alterations, including a reduction in global heterochromatin (*Kdm5a, Kdm6a, Ezh1, Ezh2, Setdb2, Smarcb1*), nucleosome remodeling (*Smarca1, Ino80b, Vps72, Phf8, Chd4, Chd5, Anp32e, Itgb3bp, Ssrp1, Baz2a*), changes in histone marks (*Mll1, Mll3, Setd2, Kdm4a, Hdac3, Hdac4, Ctcf, Cbx3*), global DNA hypomethylation, and CpG hypermethylation (*Dnmt1, Dnmt3b, Tet1, Ezh1, Ezh2*). Age-driven dysregulation of mechanisms involved in protein synthesis, folding, and turnover leads to impaired protein homeostasis, followed by the accumulation of misfolded protein aggregates. We noted the downregulation of genes involved in misfolded/unfolded protein response (*Hspb1, Hspb2, Dnajb5, Dnajb14, Dnajc24, Dnajc5, Vcpip*) and protein turnover (*Ube2e2, Usp38, Uch15, Rnf38, Fbxo31, Fbxw8*), indicating signs of impaired proteostasis.

Reduced DNA repair capability accumulates DNA damage, leading to genome instability with aging (*1*, *2*, *43*). We observed a decrease in the expression of genes involved in DNA damage response (*Trp53, Mdm2, Chek1, and Chek2*) and repair mechanisms (*Rad50, Rad52, Xpc, Xrcc1, Xrcc2, Atr, Atm, Mgmt, Apex1, Recql*), indicating diminished DNA repair in old vestibular HCs. Autophagy is crucial to recycling damaged organelles to maintain cellular homeostasis (*1*, *2*). We noted the enrichment of key genes related to autophagy initiation, autophagosome formation, and progression (*Becn1, Atg12, Atg13, Atg16l1, Atg101, Map1lc3b, Ulk1, Ulk2, Fundc1, Tbk1, Pink1, Sqstm1, Tsc2, Bnip3*). However, we also noted the downregulation of key genes (*Atg5, Atg7, Atg9a, Atg9b, Atg10, Atg14, Prkaa1, Prkaa2, Ambra1, Nrbf2, Lepr, Ppargc1a*) that are involved in autophagosome elongation, autophagy progression, and energy sensing. While this may suggest active regulation of autophagy, there are signs of dysregulation in autophagosome formation and energy sensing. Aging impacts extracellular matrix (ECM) dynamics and intercellular communication (*1*, *2*). Our data indicated a reduction in genes involved in cell-cell adhesion (*Cdh1, Cdh3, Cdh15, Pcdh17, Pcdh9, Pcdhga4, Ptprk, Ptprf*), intercellular communication (*Gja1, Gjb2, Nlgn1, Nlgn2*), ECM interactions and diminished tissue integrity (*Itgb1, Itgav, Itga8, Tjp3, Cldn10, Cldn23*).

We utilized RNAscope *in situ hy*bridization to validate some key markers identified from our analysis (Fig. 4B). Previous studies show that the downregulation of proteostasis regulator, *Hspb1* (encodes Heat-shock protein 27), is associated with ARHL (*44*). Moreover, mutations in *Gjb2* (encodes connexin 26) are associated with genetic deafness accompanied by vestibular dysfunction (*45*). Coinciding with our transcriptomic findings, we noted a significant decrease in *Hspb1* and *Gjb2* in vestibular HCs with biological aging, which may play roles in ARVD. Together, these data indicate that vestibular HC aging follows universal aging programs.

### Cell type-specific aging signatures

Previous studies have revealed distinct cell type-specific aging signatures across various tissues (*46–48*). Hence, we examined whether the universal vestibular HC aging programs are accompanied by cell type-specific aging signatures. We performed PCA multivariate analysis to examine the overall age-driven changes in the gene expression in young and old type I and II HCs across our 5 biological replicates. Our results show that the most striking differences in our datasets occur between young and old groups, rather than between type I and II HCs, as evidenced by the 96% variance of principal component 1 (Fig.5A). Next, we assessed the number of unique and shared genes expressed in young and old type I and II HCs. We noted that irrespective of cell type or age, the majority of genes (60.7%) are shared among HCs (Fig.5B). To evaluate the age-related differences between young and old HCs, we conducted a differentially expressed gene (DEG) analysis (Fig.5, C and D). We primarily focused on the genes associated with universal or cell type-specific aging signatures (Fig.5E). Our DEG analysis revealed age-related downregulation of hair bundle-related genes, including *Ush1c, Espn, Cib2, Dynll2, Tubb4b, Actb, Gsn, Calm1, Tekt2, Nudc, Evl, Cfap46,* and *Atp2b2*, most of which are associated with genetic balance disorders and hearing loss (*49–53*). Downregulation of these genes in the vestibular HCs during biological aging may play a crucial role in the development of ARVD and could serve as potential biomarkers. Additionally, many of our top enriched DEGs are linked to universal hallmarks, including oxidative stress (*mt-Co3, mt-Cytb mt-Atp6*), inflammation (*Ptgds*), and epigenetic alterations (*Pifo, Tle5, Chrac1, Gtf2i*) while top downregulated DEGs were associated with proteostasis (*Hsbp1, Hsp90ab1, Cct7, Erp29, Fbxo2, Uba52*), and intercellular communication (*Dctn3, Epcam, Afdn*). Previous studies have suggested that type I HCs are more vulnerable to aging than type II HCs (*9*). While we did not observe significant differences between the two types of HCs at the cellular and molecular levels, we observed some heterogeneity in HC function-related genes among individual HCs (fig. S3). Overall, these findings suggest that vestibular HCs exhibit both universal and cell type-specific aging signatures.

**Fig. 5.**
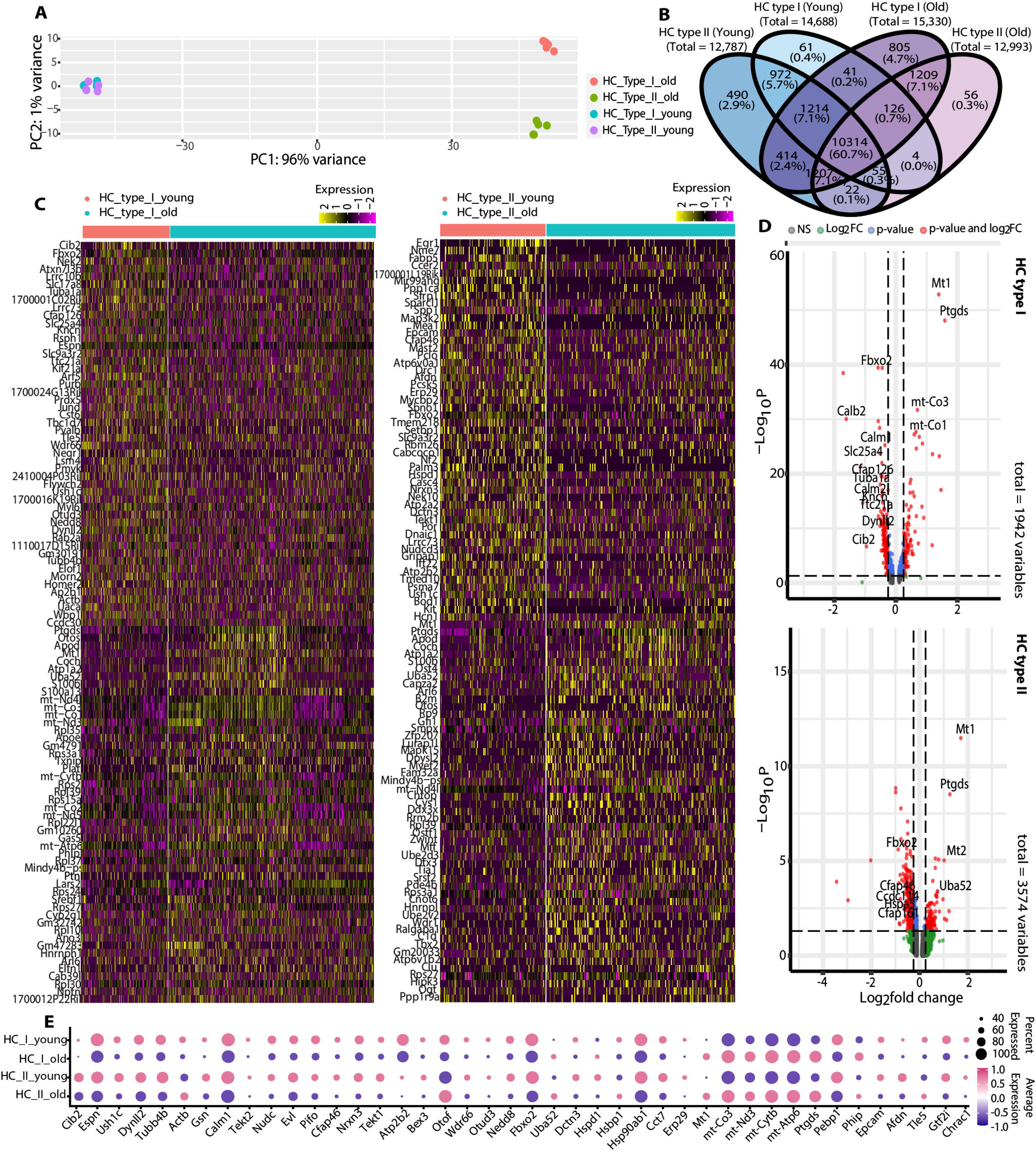
Cell type-specific aging signatures in vestibular HCs. **(A)** PCA analysis of young and old type I and II HC from the 5 biological replicates. DESeq2 in Seurat was used with the default Wald test, and multiple testing was corrected using the Benjamini-Hochberg method. Variance stabilizing transformation (VST) was applied to assess the variance among different conditions. **(B)** Venn diagram indicating the number of shared and unique genes expressed in type I and II HCs. **(C)** Heatmaps of young and old type I and II HCs indicating top 50 DEGs. Rows represent genes, columns represent single cells, with expression levels ranging from high (purple) to low (yellow), p<0.05, FCcutoff - 0.25, log2FC. threshold - 0.1. **(D)** Volcano plots of young and old type I and II HCs, p<0.05, FCcutoff - 0.25, logfc. threshold - 0.1. The x-axis represents the log2 fold- change values (avg_log2FC). The y-axis represents the raw p-values (p_val). Wilcoxon rank-sum test with Bonferroni correction for false discovery FDR<0.01 was used to analyze DEGs. **(E)** Dot plots indicating some of the key downregulated genes identified from the DEG analysis. The dot size represents the percentage of cells expressing the genes from HC type I and II clusters, whereas the color indicates the expression level. Expression levels are normalized via z-score normalization. Thus, the average expression is zero, and positive or negative values indicate expression above or below average.

### Biological processes associated with old vestibular HCs

To examine the biological relevance of our observed DEGs, we performed gene ontology (GO) enrichment analysis and Kyoto Encyclopedia of Genes and Genomes (KEGG) analysis. We observed a significant downregulation of GO terms and genes related to i) HC structure and functions including sensory perception of stimuli, stereocilia bundle, organization, and cytoskeleton, ii) proteostasis-related processes such as regulation of protein ubiquitination, proper protein folding, and refolding, iii) synapses indicating the age-driven decline of several key functional processes in old HCs (Fig.6, A and B). While kinocilia have long been regarded as a primary cilium, a recent study suggests that mouse and frog vestibular kinocilia possess both primary and motile cilia machinery, exhibiting motility (*54*). Coinciding with this, we noted downregulation of GO-terms and key genes related to kinocilia structure and motility (Fig.6, A to C). We observed a significant downregulation of many genes involved in kinocilia structure and function, such as *Ccdc39, Ccdc40, Ccdc96, Pifo, Gsn, Cfap126, Hydin, Rpsh1, Ttll3, Ttc21a, Cafp45,* as well as those linked to stereocilia and HC function such as *Espn, Ush1c, Pou4f3, Homer2, and Slc17a8.* Many of these genes are linked to ciliopathies, congenital deafness, and balance disorders (*35*, *50*, *51*, *55*).

**Fig. 6.**
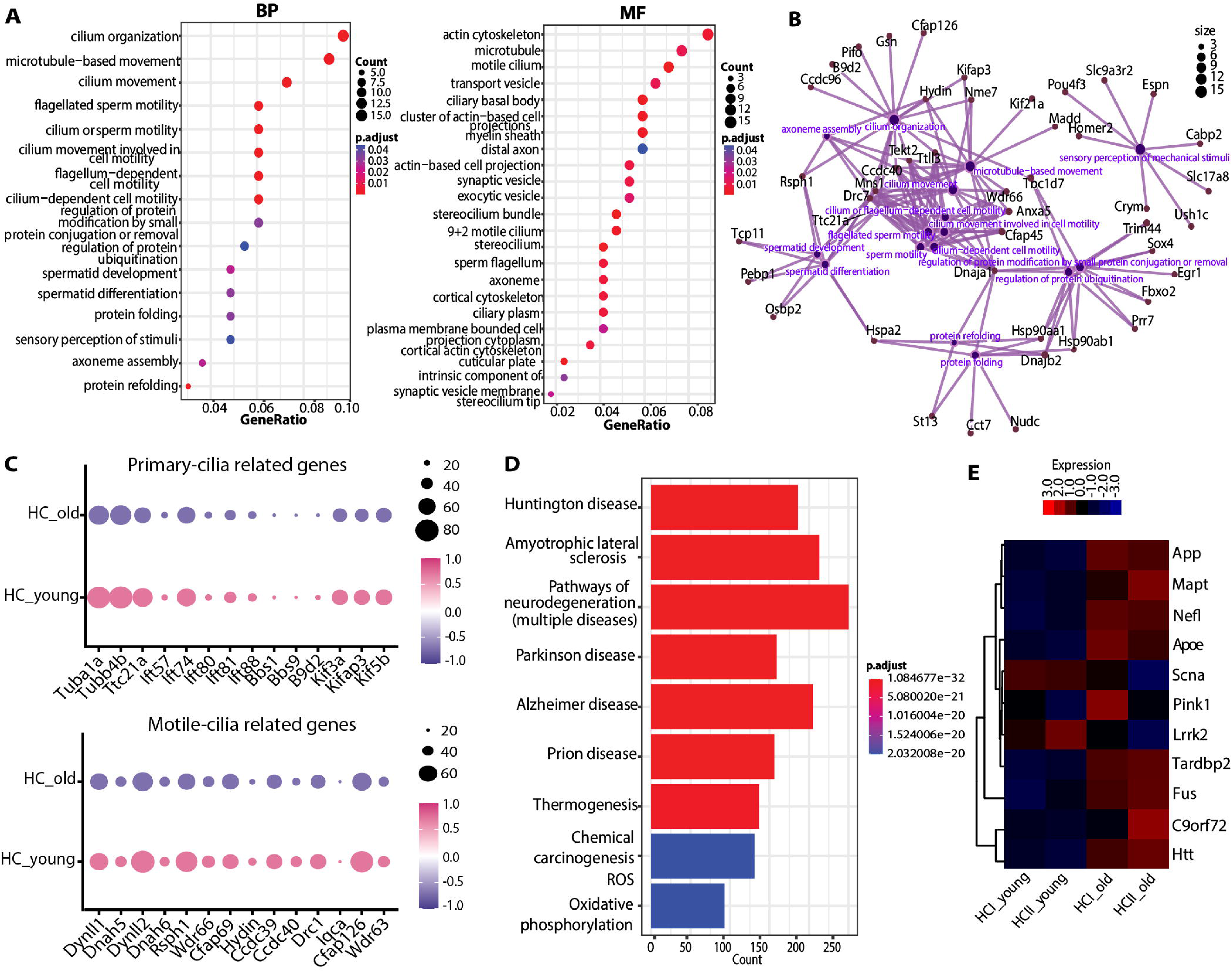
Biological processes associated with old vestibular HCs. **(A)** GO enrichment analysis of the biological process (BP) (left) and Molecular function (MF) (right). The Hypergeometric test with the Benjamini-Hochberg adjustment was used. Pairwise semantic similarity was computed for clustering the GO terms, p-value < 0.05. **(B)** Circular network (CNET) plot illustrates genes associated with the downregulated biological processes. **(C)** Dot plots indicate the genes related to kinocilia. The dot size represents the percentage of cells expressing the genes from the HC cluster, whereas the color indicates the expression level. Expression levels are normalized via z-score normalization. Thus, the average expression is zero, and positive or negative values indicate expression above or below average. **(D)** KEGG pathway analysis of old vestibular HCs. The Hypergeometric test with the Benjamini-Hochberg p adjustment was used, p-value<0.05. (**E)** Enrichment of AD, PD, HD, ALS, and dementia susceptibility genes in vestibular HCs.

ARVD is increasingly recognized as a key player in neurodegenerative diseases and cognitive decline, indicating a bidirectional association (*56*, *57*). Interestingly, our KEGG pathway analysis of old vestibular HCs revealed significant enrichment of signaling pathways linked to various neurodegenerative diseases, including Alzheimer’s (AD), Parkinson’s (PD), Huntington’s (HD), and amyotrophic lateral sclerosis (ALS), along with an elevated expression of AD/PD (*Apoe, App, Mapt*), HD (*Htt*), ALS, and dementia (*Nefl, Fus, Tardbp, C9orf72*) susceptibility genes (Fig.6, D and E) (*58–61*). Our results suggest that neurons and vestibular HCs exhibit some shared aging patterns, indicating that vestibular HC aging may represent both an associated condition and a potential biomarker for neurodegenerative diseases. These results further emphasize the complex interplay between universal and cell type-specific mechanisms underlying ARVD. Moreover, our GO analysis showed downregulation of biological processes and genes related to proteostasis, such as *Hsp90aa1, Hsp90ab1, Hspa2, Cct7*, and *Dnajb2*. Age-driven dysregulation of proteostasis and subsequent accumulation of protein aggregates lead to the onset of various diseases, including neurodegenerative diseases (*62*).

Next, we assessed the age-related downregulation of key genes identified from our analyses. Concordant with our transcriptomic findings, our experimental validations by RNAscope *in situ* hybridization and immunostaining showed a significant downregulation of key HC markers at both gene and protein levels (Fig.7, A to D, fig. S4). To examine whether these molecular changes associated with the hair bundle impair mechanotransduction in the old vestibular HCs, we assessed the FM1-43 styryl dye uptake by the vestibular HCs *in vitro*. FM1-43 is a fluorescent dye that rapidly enters HCs via mechanotransduction channels in the hair bundle (*63*). We observed a significant reduction in FM1-43 uptake in both the utricle and crista of old mice (Fig. 7E), indicating compromised mechanotransduction channel activity with aging. To further evaluate HC function *in vivo*, we recorded vestibular microphonic, which reflects HC mechanotransduction and function (*64*). We observed a significant age-associated reduction in the magnitude of vestibular microphonic response in the old mice compared to young mice (Fig. 7F), supporting the conclusion that aging compromises both mechanotransduction and overall HC performance.

**Fig. 7.**
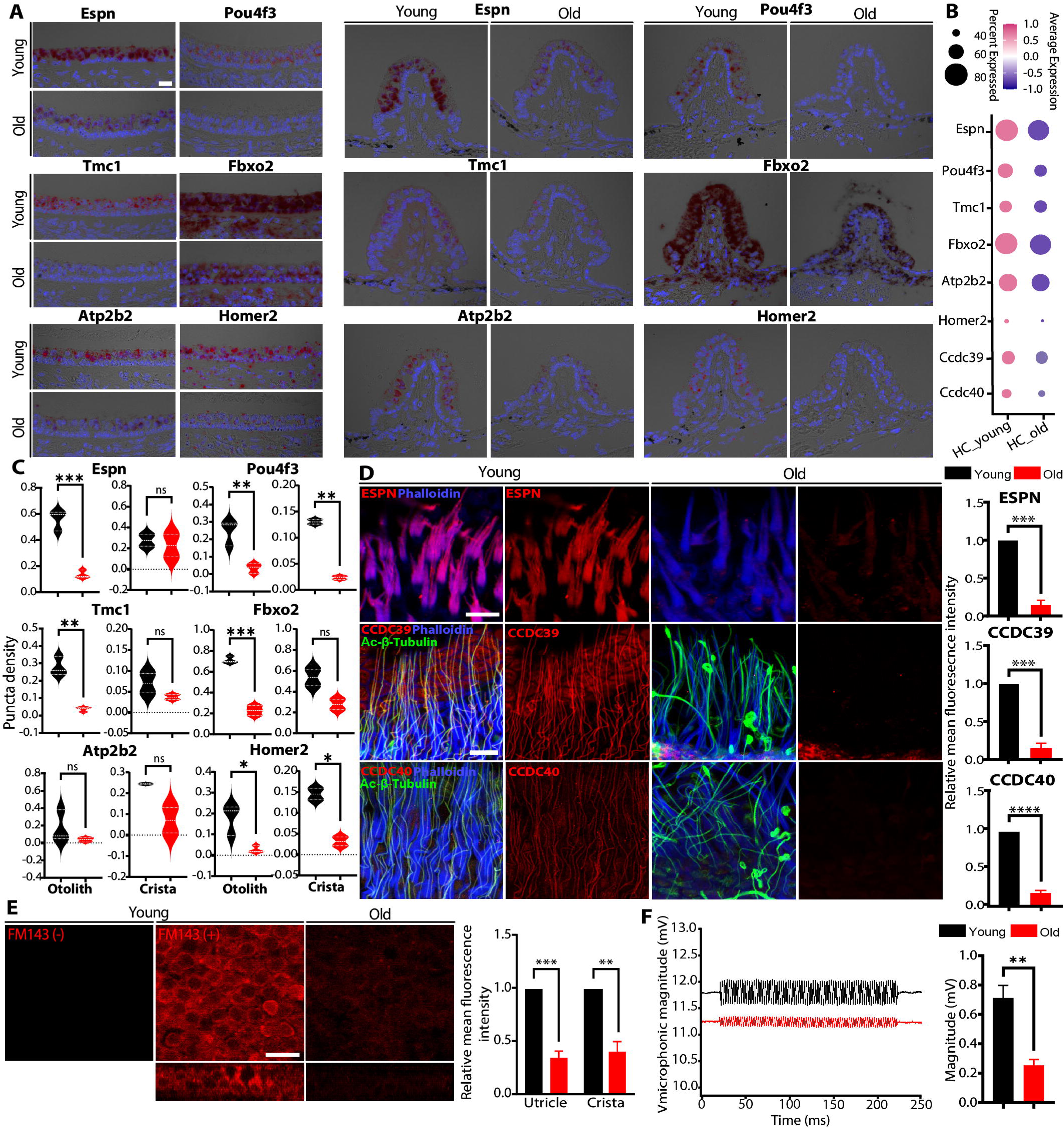
Age-related reduction of key hair bundle markers and HC function. **(A)** RNAscope *in situ* hybridization validations of key HC genes identified from DEG and GO analysis in young and old otoliths and crista. Scale bar, 10 µm. **(B)** Dot plot shows the average gene expression levels in young and old HCs. **(C)** Quantifications of RNAscope *in situ* hybridization from otoliths (n=3) and crista (n=2). Data shown as mean ± SEM, ns-non-significant, *p <0.05, **p <0.01, ***p <0.001 by unpaired t-test. **(D)** Representative images depicting age-related reduction in stereocilia-related (ESPN) and kinocilia-related (CCDC39, CCDC40) markers (red) and quantifications of relative mean fluorescence intensity (n=3). Scale bar, 10 µm. Data are shown as mean ± SEM, ***p<0.001, ****p<0.0001 by unpaired t-test. **(E)** Representative images (utricle) and quantification of relative mean fluorescence intensity of FM1-43 dye uptake in young and old HCs in the utricle and cristae (n=3). Scale bar, 10 µm. Data are shown as mean ± SEM, **p <0.01, ***p <0.001 by unpaired t-test. **(F)** Representative images of vestibular microphonic measurements from young and old macula of the utricle (n=5 per age group). The stimulus was a 390 Hz sinusoid signal from a Burleigh PZ-150M Driver.

### Tissue-specific age-related changes in vestibular and cochlear HCs

The cochlea and vestibule display a different pace of aging, leading to a differential onset and progression of ARHL and ARVD. However, the cellular and molecular processes underlying this difference remain poorly understood (*9*, *65*). We measured auditory brainstem response (ABR), distortion product otoacoustic emissions (DPOAE), and endocochlear potential (EP) to investigate auditory functional decline in our animal cohorts in addition to VsEP (fig. S5A-C). In line with previous studies, we observed a significant elevation in ABR and DPOAE thresholds during aging (*66*). Although some studies from gerbils have reported an age-related decline in EP, our findings revealed no changes in EP in mice. Next, we assessed the age-related morphological changes in the cochlea. The cochlea contains inner and outer HCs (IHCs and OHCs), which are organized tonotopically along the length of the cochlea. Previous findings indicate that the apical and basal turns of the cochlea, which are responsible for detecting low and high frequencies, respectively, show greater vulnerability to aging (*33*, *67*). We observed a significant HC loss, especially in the apical and basal ends of the cochlea (fig. S5D). We also noted age-related hypertrophy and atrophy in the cochlear HCs. In particular, we observed atrophy of both IHCs and OHCs in the apical turn, and OHC atrophy and IHC hypertrophy in the basal turn (fig. S5E).

Next, we assessed age-associated tissue-specific changes between vestibular and cochlear HCs at the molecular level. We conducted a GO analysis to examine the biological processes downregulated in old cochlear HCs compared to age-matched vestibular HCs. We noted a significant downregulation of HC-specific functions, including sensory perception and response, stereocilia-related processes, and synapses in cochlear HCs. We also observed the downregulation of universal processes such as oxidative stress, DNA damage, inflammatory response, and diminished proteostasis and autophagy compared to age-matched old vestibular HCs (Fig.8A and fig. S6). Concordant with that, we observed the age-related downregulation of key genes essential for HC function in old cochlear HCs (Fig.8B). Comparison of age-related log2 fold changes in HC-related gene expression in cochlear and vestibular HCs revealed more pronounced downregulation in cochlear HCs compared to vestibular HCs (Fig.8C). Accumulation of various degenerative molecular changes with aging trigger cellular senescence. It is typically characterized by increased senescence-associated-β-galactosidase (SA-β-gal) activity at pH 6.0 (distinguishing from general β-gal at pH 4.0), allowing selective staining of senescent cells by cleaving X-gal (5-bromo-4-chloro-3-indolyl-β-D-galactopyranoside), resulting in a blue precipitate (*68*, *69*). A previous study shows increased SASP and SA-

**Fig. 8.**
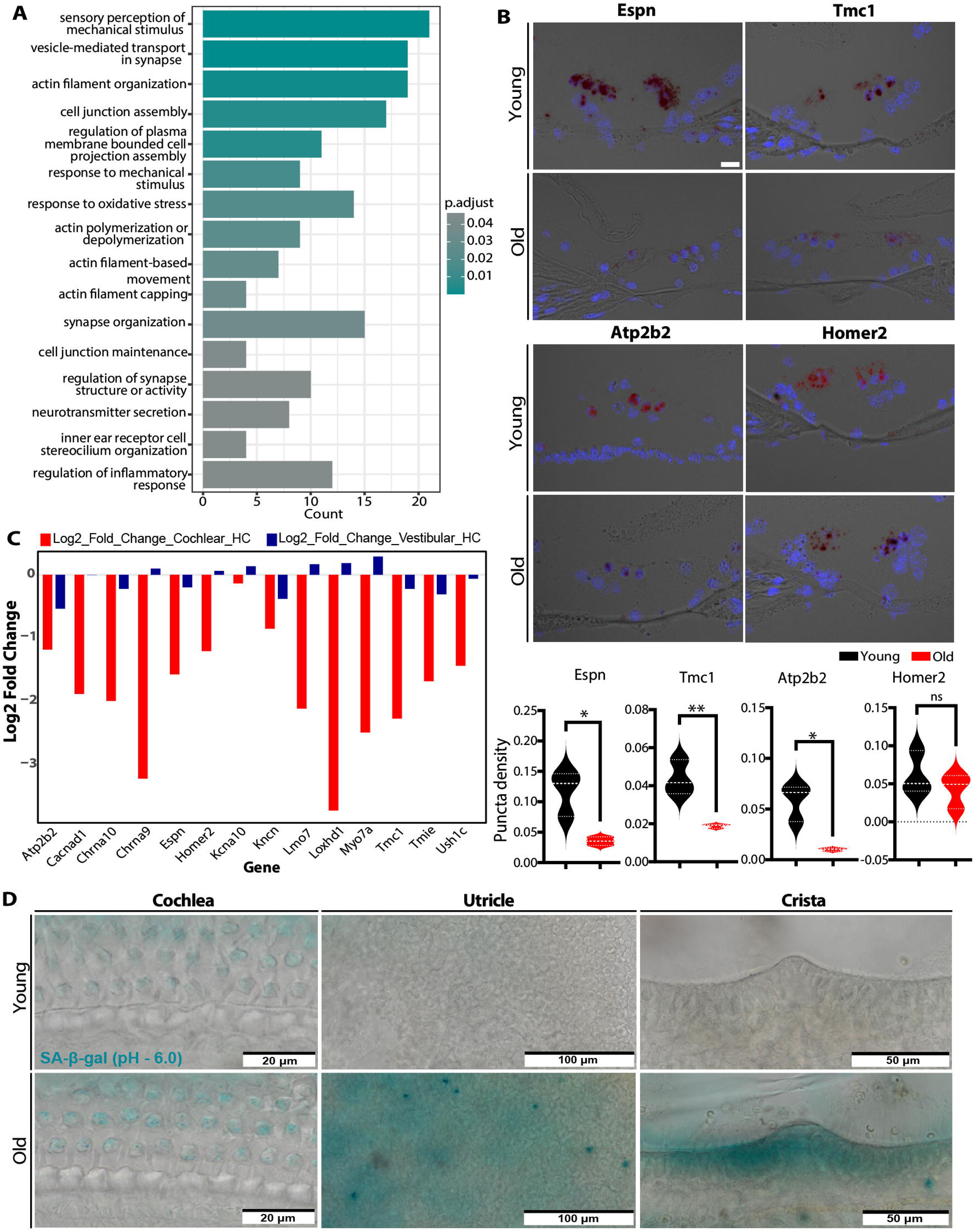
Age-related changes in cochlear and vestibular HCs. **(A)** GO analysis showing biological processes downregulated in the old cochlear HCs compared to old vestibular HCs. The top genes that were downregulated in old cochlear HCs compared to age-matched old vestibular HCs were (Log2FC<0) used for this analysis using the Hypergeometric test with the Benjamini-Hochberg p adjustment, p<0.05. Pairwise semantic similarity was computed to cluster the GO terms. **(B)** RNAscope *in situ* hybridization validations and quantifications in cochlear HCs (n=3). Scale bar, 10 μm. Puncta density was quantified for each gene using FIJI (n=3). Data shown as mean ± SEM, ns-non-significant, *p <0.05, **p <0.01 by unpaired t-test. **(C)** Age-related Log2 fold changes of key genes related to HC functions in cochlear and vestibular HCs. **(D)** Senescence-associated β-galactosidase (SA-β-gal) activity at pH 6.0 in the cochlear and vestibular sensory epithelia with aging, indicating cellular senescence.

β-gal activity in the cochlear HCs upon hydrogen peroxide (H_2_O_2_)-induced DNA damage in an accelerated aging mouse model (*70*). Thus, we assessed SA-β-gal activity in the cochlear and vestibular HCs, and we observed an increase in SA-β-gal activity in both old cochlear and vestibular HCs (Fig.8D). Unlike young vestibular HCs, we also noted the expression of SA-β-gal in young cochlear HCs, indicating potential early onset of age-related molecular changes in cochlear HCs. These findings highlight subtle tissue-specific differences in the aging processes between vestibular and cochlear HCs.

### Aging signatures in human vestibular HCs

Previous studies have reported various degenerative changes in vestibular HCs in elderly individuals with ARVD (fig. S7A) (*71*). The age-related changes of the vestibular end organs begin around the age of 55, and more drastic deterioration, including HC loss, occurs after 70-80 years in humans (*9*). However, the molecular changes may occur earlier than the functional and structural changes, gradually accumulating damage and compromising HC integrity. Understanding the age-related molecular changes of the human vestibular HCs remains a significant challenge due to the limited availability of human samples. To examine the aging signatures in human vestibular HCs, we utilized two publicly available transcriptomic data sets from human organ donors (GSE213796) (GSE207817) (*72*, *73*).

We analyzed vestibular HC datasets from 9.2 fetal weeks and 45-year-old donors for comparison. Our analysis focused on universal hallmarks, as developmental specializations of HCs may still be ongoing at the age of 9.2 fetal weeks. Moreover, 45-year-old datasets were acquired by single-cell RNA sequencing, whereas the fetal data were acquired by single-nucleus RNA sequencing, which excludes cytoplasmic transcripts such as mitochondrial genes. Consequently, we excluded some pathways, such as oxidative stress, from the analysis due to the differences in transcriptomic data acquisition methods. Data were pre-processed and clustered using standard transcriptomic analysis pipeline, and HC clusters were identified using known marker genes (fig. S8). Raw count data from the HCs were extracted and aggregated by averaging counts to generate a pseudobulk data frame containing summed expression values. These values were then scaled and log2-normalized to assess the age-related changes (fig. S7). We observed a reduction in expression of genes linked to global heterochromatin and nucleosome remodeling, changes in histone marks, global DNA hypomethylation, and CpG hypermethylation, indicating epigenetic alterations. Additionally, we noted a substantial downregulation of genes related to DNA damage repair mechanisms, ECM, and intercellular communication. While key genes related to autophagy initiation and progression were enriched, some key genes involved in autophagosome formation were downregulated.

## Discussion

Despite the high prevalence of ARVD, its molecular basis remains inadequately understood, hindering the development of targeted treatments (*5*, *9*). In this study, we conducted a comprehensive transcriptomic analysis alongside functional and morphological assessments to examine age-related alterations in the vestibular end organs in mice. We identified genes and molecular processes associated with both universal and cell type-specific aging signatures in the vestibular HCs. Additionally, we uncovered tissue-specific molecular differences in aging between cochlear and vestibular HCs, providing insights into differences in age-related processes within these two mechanosensory systems.

Aging cells, including post-mitotic neurons, photoreceptor cells, and cochlear HCs, exhibit universal aging hallmarks (*33*, *46*, *74*), and we demonstrated that vestibular HCs also conform to these hallmarks. Our data indicated an increase in senescence, marked by a decrease in cyclin-dependent kinases (CDKs), and an increase in cyclin-dependent kinase inhibitors (CDKis) alongside alterations in SASP genes related to laminins, heterochromatic foci, apoptosis, hypoxia, and angiogenesis in old HCs (Fig.4). This was further evident by elevated SA-β-gal staining in old vestibular HCs (Fig.8). Oxidative stress and inflammation play a critical role in ARHL (*75*). Similarly, we observed signatures of increased oxidative stress and low-grade inflammation in old vestibular HCs. Mitochondrial electron transport chain (ETC) byproducts are a major contributor to ROS generation. We detected an age-related increase in the expression of multiple ETC genes, accompanied by elevated expression of antioxidant genes, suggesting increased ROS production along with compensatory clearance. However, we noted that genes related to H_2_O_2_ free radical clearance were downregulated, suggesting partial ROS clearance. Additionally, genes involved in mitochondrial stress response and quality control were upregulated, further supporting the presence of increased oxidative stress and the activation of adaptive protective mechanisms. While our findings suggested active maintenance of autophagy, decreased expression of several genes involved in autophagosome formation and energy sensing indicate signs of impaired autophagy.

We observed an age-related decrease in genes involved in DNA repair mechanisms, indicating diminished DNA damage repair efficiency in old vestibular HCs in mice and humans. Moreover, we noted significant epigenetic alterations with aging, including alterations in histone marks, DNA methylation, global heterochromatin, and nucleosome remodeling. We also noted a significant reduction in genes involved in ECM, intercellular adhesion, and communication, including cadherins, protocadherins, and integrins, in the old vestibular HCs. Age-related downregulation of these genes may contribute to the degeneration of tip/lateral links, leading to stereocilia and kinocilia fusion (Fig.2). These genes also play a crucial role in cell adhesion, and reduction may lead to an age-related decrease in cell-cell adhesion. Prior studies show a correlation between neurodegenerative diseases and ARHL and ARVD (*57*). Dysregulated proteostasis plays a crucial role in the accumulation of harmful protein aggregates, leading to neurodegenerative diseases (*62*). Overexpression of β-Amyloid and tau proteins in cochlear HCs of Alzheimer’s mouse model indicated severe hearing defects and significant HC loss with aging (*76*). Interestingly, we observed the enrichment of neurodegenerative pathways and increased expression of neurodegenerative susceptibility genes, including *Mapt* (encodes for tau) and *App* (encodes for amyloid-β precursor protein), in old vestibular HCs (Fig.6), suggesting shared aging patterns between neurons and vestibular HCs.

Due to the multifaceted decline of oculomotor, postural, perceptual, and cognitive functions in ARVD, age-related synaptic and neuronal loss has been considered a contributor to ARVD. While there are mixed findings on age-related synaptic and vestibular ganglion neuron density, recent reports have shown no significant loss of synapses (*3*) and vestibular neurons (*77*). Our findings indicated no substantial changes in genes related to key Ca^2+^ and K^+^ channels (fig. S3). Strikingly, the most prominent age-related changes were observed in the genes associated with the hair bundles and mechanotransduction apparatus (Fig. 5 to 7), which is highly consistent with the age-related stereociliopathy and kinocilia degeneration observed at the morphological level (Fig.2). Interestingly, most of the down-regulated genes, including *Ush1c, Tmc1, Pou4f3, Espn*, *Fbxo2, Atp2b2, Slc17a8, Homer2, Ccdc39, Ccdc40,* and *Tekt2*, are associated with congenital hearing and balance disorders (*49–51*, *55*). Based on the wear and tear theory of aging (*78*), hair bundles are particularly vulnerable to aging as they are the first in line to respond to mechanical stimuli, causing constant mechanoexposure. Since no significant vestibular HC loss was observed at 24 months (Fig.1), the age-related vestibular functional decline may not result from HC loss, but rather from degeneration of the hair bundle structure and mechanotransduction machinery, leading to ARVD (Fig. 7).

Numerous studies have shown diverse molecular changes in various sensory organs during biological aging. These changes vary based on the inherent differences in cellular dynamics in each organ, which regulate the degree and pace of aging (*47*, *48*). Examination of the aging signatures of old cochlear HCs relative to age-matched vestibular HCs indicated significant downregulation of processes and genes related to cell type-specific (stereocilia, mechanotransduction) and universal programs (Fig. 8 and fig. S6), aligning with bundle degeneration, cellular hypertrophy/atrophy, and increased loss of cochlear HCs (fig. S5) (*33*). We also noted an early onset of cellular senescence in cochlear HCs compared to vestibular HCs, as evidenced by SA-β-gal activity in young cochlear HCs (Fig.8). This indicates a relatively early onset of aging in cochlear HCs compared to vestibular HCs. However, it remains unclear whether these differences reflect distinct molecular underpinnings. The divergence in aging onset may be influenced by multiple factors, including extrinsic influences (e.g., noise exposure, ototoxic drugs, lifestyle, and systemic health conditions) and intrinsic factors (e.g., genetic predisposition). Moreover, it is also unclear whether the limited capability of vestibular supporting cells to HC conversion may also influence this process.

This study has some limitations. Our findings warrant further investigation, testing mechanistic hypotheses, and using genetic knockout models to examine how these genes impact vestibular function. We assessed the age-related changes in HCs by pooling the HCs from the three vestibular end organs together, which may mask important regional differences in age-related molecular changes. Although we attempted to characterize aging signatures in the human vestibular HCs, these analyses require validation across appropriate age groups to capture both cell type-specific and universal aging signatures. While our findings provide some insights into differences in age-related changes in cochlear and vestibular HCs, it remains challenging to pinpoint a single principle mechanism underlying the difference in the pace of aging between vestibular and auditory organs.

To the best of our knowledge, this is the first study to comprehensively characterize both universal and cell type-specific aging signatures of the vestibular HCs, providing insights into previously unrecognized aspects of vestibular aging. Both universal and cell type-specific aging processes at the molecular level may begin early, leading to the gradual disruption of function, loss of homeostasis, and degenerative changes in vestibular HCs preceding the functional and morphological decline observed at later stages. While our study highlights bundle degeneration and the associated decline in mechanotransduction as a key contributor, it also provides a comprehensive understanding of the underlying mechanisms and molecular drivers of ARVD, paving the way to develop targeted treatment strategies to mitigate ARVD.

## Methods

### Experimental animals

Wild-type male and female CBA/J mice used in this study were purchased from the Jackson Laboratory (Stock #:000656) and were bred and aged in the Animal Facility of Creighton University under standard conditions. The acoustic environment of the animals was examined by a sound level meter (CEL-500, Casella Cel, UK). The sound levels were 68 dB SPL over 98% of the time, and 75 dB SPL 1-2% of the time. All experimental procedures were approved by the Institutional Animal Care and Use Committee of Creighton University.

### Vestibular Sensory Evoked Potential (VsEP)

Animal preparation and VsEP testing were performed as previously described (*79*). Briefly, mice were anesthetized with a combination of Ketamine (126 mg/kg)/Xylazine (14 mg/kg) injected intraperitoneally and supplemented as needed to maintain the anesthesia. Body temperature was maintained at 37±1°C via a homeothermic heating blanket. The recording electrodes were placed-anterior to the nuchal crest (non-inverting), behind the pinna (inverting), and over the hip (ground) subcutaneously. A custom head clip was used to secure the head to the mechanical shaker (ET-132 Labworks Inc.). The head translation was induced along the naso-occipital axis at a rate of 17 linear pulses per second with alternating polarity and stimulus intensities ranging from +6 to -18 dB relative to 1.0 g/ms (1.0 g = 9.8 m/s^2^) adjusted in 3 dB intervals. A broad-band forward masker (50-50,000 Hz, 92 dB SPL) was presented to avoid auditory responses. 128 responses from each of the two polarities (upward and downward head translations) were averaged to yield the mean VsEP response. First positive (p1) and negative (n1) response peaks correspond to the compound action potential from the vestibular nerve. Thus, the amplitude and latency of these peaks were quantified. The threshold was defined as the mid-level of the lowest stimulus intensity that elicited a detectable response and the next lowest level that did not produce a detectable response.

### Auditory brainstem response (ABR) and Distortion Product otoacoustic emissions (DPOAE)

ABRs were recorded from young and old mice in a soundproof chamber as previously described (*33*). The mice were anesthetized with a mixture of ketamine/xylazine, and the body temperature was maintained using a heating pad as mentioned in the VsEP procedure. Platinum recording electrodes were placed at the vertex (non-inverting), mastoid prominence (inverting), and leg (ground) subcutaneously. Tone bursts ranging from 4-50 kHz were used as the stimuli, and ABRs were amplified (100,000x), filtered, and recorded by the TDT RZ6 (Tucker-Davis Technologies, Alachua, FL). 200 stimulus repetitions were used to yield the averaged responses. The lowest sound pressure level (dB) at which any wave (wave I to IV) was visibly detected and reproducible above the noise level was regarded as the ABR threshold. The DPOAE at 2f1-2f2 was recorded in response to f1 and f2, with f1/f2=1.2 and the f2 level was set 10 dB lower than f1. The sound signal was recorded from the inner ear canal via a microphone, amplified, and the fast Fourier transforms were computed from the averaged waveforms obtained from the inner ear sound signal. The f1 sound pressure required to produce a response above the noise level at the frequency of 2f1-f2 was considered the DPOAE threshold. Recorded ABR and DPOAE thresholds were graphed and statistically analyzed using GraphPad Prism.

### Endolymphatic potential (ELP) and endocochlear potential (EP)

ELP and EP measurements from the young and old mice were performed as described previously (*27*). Post anesthesia, tracheotomy was performed in the ventral position without providing artificial respiration. Tissue and musculature overlying the tympanic bulla were removed, and the bulla was opened to access the cochlea. A fine drill was used to make small holes in the lateral wall in the apical and the basal turns of the cochlea. A glass capillary microelectrode (5-8 MΩ) filled with 150 mM KCl was mounted on the Leica micromanipulator. A stable positive DC potential was observed when microelectrode entered the scala media. The responses were amplified (high-pass filter at 1 kHz) using an Axopatch 200B amplifier (Molecular Probe, Sunnyvale, CA, USA) under current-clamp mode and acquired by software pClamp 10 running on an IBM-compatible computer with 16-bit A/D converter (Digidata 1440A). The voltage changes during the penetration were recorded under the gap-free mode via Clampex in the pClamp software (version 10, Molecular Devices) with a sampling frequency of 1 kHz. ELP in the utricle was recorded by inserting the microelectrode through the round window, advancing across scala media, and scala vestibuli to the utricle. The tip of the microelectrode was loaded with fluorescent phalloidin (Invitrogen #565227) diluted in 150 mM KCl (1:100) and injected into the utricle to confirm that the recording was made from the utricle. Post-recording utricle was processed and observed under a confocal microscope.

### Recording of vestibular microphonic from utricle maculae

The vestibular microphonic was recorded as described previously (*64*). Vestibular microphonic from the macula of utricle was evoked by driving the inner ear fluid (ref). The stimulus was a 390 Hz sinusoid with a duration of 300 ms and a rise-time of 1 ms from a Burleigh PZ-150M Driver. The piezo actuator was attached to the stapes foot through a tapered glass rod (1.5 cm in length and 0.8 mm in diameter at the stapes foot) after the ossicular chain was severed. The peak-to-peak magnitude of the piezo motion was preset at 3 μm, sufficient to drive a saturated response. We did not expect the contribution of cochlear microphonic to our recording, as the frequency range of the basilar membrane response of CBA/J is above 2 kHz. To rule out the possibility of contribution by cochlear microphonic, the helicotrema in the apical turn of the cochlea was destroyed before the vestibular microphonic was recorded. The microphonic signal from the microelectrode was amplified and high-pass filtered (at 1 kHz) under current-clamp mode using an Axopatch 200B amplifier and acquired by software pClamp 10 running on an IBM-compatible computer with a 16-bit A/D converter (Digidata 1440A). The sampling frequency was 10 kHz. Response from 20 presentations was averaged for each recording.

### Scanning electron microscopy (SEM)

Utricle and cristae from the two age groups were dissected and fixed in 2% glutaraldehyde, and 4% PFA in 0.1M sodium cacodylate buffer (pH 7.4) overnight. The fixed samples were washed with 0.1 M phosphate buffer. After removing the otolithic membrane from the utricle, the samples were treated with 1% Osmium tetroxide and washed with 0.1M sodium cacodylate buffer. The samples were then dehydrated using a series of ethanol, followed by critical point drying using carbon dioxide via critical point dryer (EMS 850) and mounted onto the SEM stubs. The morphology of the stereocilia bundle was visualized, and the images were acquired via FEI Quanta 200 scanning electron microscope (ThermoFisher, Hillsboro, OR).

### Cell dissociation, cDNA library preparation, and single-cell RNA sequencing

Male and female mice aged 2.5 months (young) and 22 months (old) were used for scRNA-seq. Whole cochlear and vestibular organs were microdissected from the inner ears of CBA/J mice in petri dishes containing cold L-15 media (Gibco #11320033). The sensory epithelia were subjected to enzymatic digestion by transferring them into 1.5 ml eppendorf tubes containing 1 ml/mg collagenase IV (Sigma) in L-15 medium for 10 minutes at room temperature. Then the cells were resuspended in 400 µL of enzyme-free DMEM containing 10% fetal bovine serum. Single-cell suspensions were obtained by trituration using a trimmed 200 µl pipette tip and filtered with a 40 µM strainer. The cells were pelleted by centrifuging at 300 xg for 5 mins at 4°C, then resuspended in media and subsequently quantified the number of cells. 7-8 mice were used for each biological replicate, and 5 biological replicates were obtained for the cochlea and vestibule for the two age groups, respectively. Single-cell capture, and library preparation were performed using droplet-based microfluidic technology, following the manufacturer’s instructions for the Chromium Next GEM Single Cell 3ʹ v3.1 dual index kit. The quality of the obtained libraries was assessed via high-sensitivity DNA kits (Agilent Technologies) using Agilent 4200 TapeStation prior to sequencing with Illumina NovaSeq 6000, acquiring 50,000 reads per cell.

### Raw data processing and quality control

Standard 10x Genomics CellRanger pipeline (version 6.1.2) was used for demultiplexing, alignment (10x mouse reference genome mm10), barcode processing, and UMI counting to obtain the filtered count matrices. Processing and visualization of the scRNA-seq data from cochlea and vestibule were performed in R (version 4.2.3). Ten filtered count matrices obtained from 10 biological replicates (5 per age group) from cochlear and vestibular samples were loaded into R, converted to Seurat Objects, and pre-processed to eliminate the poor-quality cells using parameters such as counts, features, percentage of mitochondria, ribosomes, and blood gene content. Cells with at least 200 features, 1000 UMI counts, and less than 25% mitochondrial genes were included for the downstream analysis. Individual Seurat Objects were integrated into a single Seurat Object for each age group, resulting in two main Seurat Objects for young and old samples in both cochlea and vestibule, respectively, creating four Seurat Objects in total.

### Normalization, scaling, integration, dimensional reduction, and cluster annotation

Integrated objects were then normalized using log-normalization and scaled using the ‘NormalizeData’ and ‘ScaleData’ functions, followed by unsupervised principal component analysis (PCA) and dimensional reduction. The first 30 PCs were used for the dimensional reduction. The ‘FindNeighbors’ was used to calculate pairwise similarities and the ‘FindClusters’ function, which uses the Louvain algorithm, was applied with the optimal cluster resolution for all four Seurat Objects, respectively. The top markers of each cluster were identified using the ‘Findmarkers’ function. Samples were then visualized via uniform manifold approximation and projection (UMAP). This dimensional reduction method uses k-nearest neighbors and Euclidean distance algorithms, preserving the local and global structure of the data. A total of 33 and 27 distinct clusters were identified from the initial clustering of young and old vestibular samples, while 31 and 33 clusters were identified from young and old cochlear samples, respectively. Previously reported cell-type-specific pan markers were used to classify the cell-type distribution in young and old vestibular and cochlear samples. Based on the known marker genes, we identified many cell types, including vestibular type I and II HCs as well as cochlear IHCs and OHCs. Vestibular and cochlear HC clusters from the two age groups were then subsetted out and reclustered for the downstream analysis to investigate age-related molecular changes.

### Pseudobulk gene expression analysis

Subsetted out young and old type I and II HCs were used to extract raw count data from single cells and aggregated them by averaging counts across young and old biological replicates to generate a pseudobulk data frame that contains summed expression values for the 4 HC groups (young and old type I and II). Expression values were then scaled and log2-normalized (table S2). Genes associated with universal aging pathways were gathered manually from multiple publicly available resources such as Aging Atlas, GenAge, SASP Atlas, and literature. Then, the general trends in the age-related expression changes were evaluated using genes associated with each aging pathway using the ‘ComplexHeatMap’ function. Expression data were extracted and centered using Cluster 3.0 (version 1.59), and heatmaps were generated using JAVA TreeView (version 1.2.1). For human vestibular HC analysis, the data sets were obtained from the GEO, pre-processed, and annotated the clusters following the same pipeline used for the murine pseudobulk analysis.

### Differentially expressed gene (DEG) analysis

Principle component analysis (PCA) was performed to assess the variance among young and old vestibular datasets by using the ‘DESeq2’ package in Seurat. The default statistical test in the ‘DESeq2’ package is the Wald test with Benjamini-Hochberg correction for multiple testing. DESeq2 uses a generalized linear model (GLM) to identify DEGs within multiple groups, and variance stabilizing transformation (VST) is used to assess the variance, revealing the biological patterns in data. DESeq2 pseudo bulk differential expression pipeline was used to generate Venn diagram via ggplot2 based ‘ggVennDiagram’ package. The Wilcoxon rank-sum test with Bonferroni correction in Seurat was used to assess the differentially expressed genes in young and old samples and subsequently visualized as heatmaps and volcano plots. The top 50 DEGs were identified using the ‘FindMarkers’ function, using young HCs as the reference and old HCs as the test group to generate heatmaps using the ‘DoHeatmap’ function. The genes at least expressed in 25% of cells in the cluster were considered for this analysis and ordered according to Log2 fold change (Log2FC). The ‘EnhancedVolcano’ function was used to generate the volcano plots using a fold-change cut-off (FCcutoff) value of 0.25 and a p-value (pCutoff) of 0.05.

### Functional enrichment analysis

The biological relevance of the top DEGs in the old samples compared to the young samples was assessed using gene ontology (GO) enrichment and over-representation analysis. EnrichGO in the ‘clusterProfiler’ package in Seurat was used to conduct this analysis. This algorithm uses the Hypergeometric test with the Benjamini-Hochberg method for multiple testing corrections. The top genes that were downregulated with aging (Log2FC<0) were used to assess the biological processes (BP), and molecular function (MF). Pairwise semantic similarity was computed for clustering the GO terms, and data were visualized as dot plots, emap plots, and cnet plots. Kyoto Encyclopedia of Genes and Genomes (KEGG) analysis was performed to assess the pathways enriched in old vestibular HCs (Log2FC<0). DESeq2 was used to generate a pseudo bulk data matrix as described above in the DEG analysis. The ‘enrichKEGG’ function in the ‘clusterProfiler’ package was used for this analysis. This algorithm also uses the Hypergeometric test with the Benjamini-Hochberg adjustment.

### Histology

Temporal bones were isolated from the young, and old mice, and a hole was made in the apical cochlea followed by overnight fixation with 4% PFA (Electron Microscopy Science, Cat: 15710) in PBS (Cytvia, SH30258.02). Tissues were thoroughly washed with 1x PBS, cleared with xylene, and rehydrated with a series of ethanol. Then tissues were embedded in paraffin wax, and the 8-10 µm thick sections were made using a microtome. H&E staining was performed using Fisher Scientific Gemini^TM^ AS automated slide stainer. The slides were loaded onto the Gemini AS basket and immersed in xylene for 3 minutes (x3), followed by a series of ethanol washes (100%, 95%, and 70%), 2 mins each. Next, the slides were immersed in hematoxylin for 2 mins, rinsed with water, and clarifier two was added (20 seconds) followed by 1 min rinse with water. Bluing reagent was added for 1 min, followed by 1 min water rinse. Slides were then submerged in 70% and 95% ethanol (2 mins each). Then the slides were immersed with counterstain eosin for 1 min and the slides were then immersed in 100% ethanol (2 mins X3) followed by incubation with xylene (2 mins x3) and mounting. Images were taken using Olympus VS120 virtual slide scanner.

### Immunostaining and Confocal imaging

Temporal bones were isolated from the young, and old mice, and a hole was made in the apical cochlea followed by overnight fixation with 4% PFA (Electron Microscopy Science, Cat: 15710) in PBS (Cytvia, SH30258.02) solution at 4°C. For the whole mount preparations, the vestibule and cochlea were microdissected from the temporal bones, then permeabilized in 1x PBST (0.3% Triton-X-100 in PBS) for 30 min followed by blocking with 10% normal goat serum (Sera care, Cat: 5560-0007) for 1h at room temperature. Tissues were incubated with primary antibodies overnight at 4°C on a rocker. The samples were thoroughly washed with 1x PBST for 30 min and incubated for 40 min at room temperature with secondary antibodies and counterstains such as DAPI and Phalloidin. The tissues were subsequently washed with 1x PBST for 40 min and carefully mounted with Fluoromount mounting media. For cryosections, the tissues were fixed overnight with 4% PFA in PBS solution at 4°C and decalcified in 10% EDTA (E671001-0500; Sangon Biotech). Tissues were washed with PBS post-fixing and cryoprotected in a sucrose gradient of 15% and 30%. The sections were then embedded in OCT (Fisher Healthcare, Cat: 4585) and kept on dry ice. 10 μm thick sections were made using Cryostat (Leica CM 1950). We used primary antibodies to immunolabel the following markers: anti-ESPN, anti-CCDC39, anti-CCDC40, anti-MYO6, anti-MYO7A, anti-acetylated-β-Tubulin, anti-SOX2, Phalloidin Flour^TM^ 488, Phalloidin Flour^TM^ Plus 405. Goat anti-rabbit Fluor 568, donkey anti-mouse 568, goat anti-rabbit Fluor 488, goat anti-mouse Fluor 488 secondary antibodies were used (table S1). Confocal images were taken via Zeiss LSM 700 upright microscope, Zeiss LSM 700 inverted microscope, and Zeiss LSM 980 inverted confocal microscope. Optical image sections were captured using orthoslicer of Imaris application, and Adobe Photoshop was used to remove the background and isolate individual hair cell images.

### RNAscope *in situ* hybridization

Decalcification and microdissection of young and old vestibular and cochlear tissues were performed as described above in the immunohistochemistry. Gene expression validations were performed using RNAscope *in situ* hybridization following the manufacturer’s instructions for RNAscope^TM^ 2.5 HD Red Assay from Advanced Cell Diagnostics (ACD Bio, Cat: 322360). In brief, paraffin-embedded tissue sections of 10 µm were baked in the HybEZ oven at 60°C followed by 10 mins incubation with xylene and ethanol to dewax and dehydrate, respectively. Sections were then subjected to hydrogen peroxide and protease treatment before incubation with probes for 2h at 40°C. Subsequently, amplification, detection, and counterstaining with DAPI were carried out before the samples were mounted. Images were acquired via Zeiss 700 inverted confocal microscope, and the obtained z-stacks were presented in maximum-intensity projection. Following this procedure, 10 probes were used to assess the gene expression, including *Espn*, *Tmc1*, *Atp2b2*, Fbxo2, *Pou4f3*, *Homer2*, *Gjb2*, *Hspb1*, and *DapB* (negative control). The quantification of RNAscope puncta density was performed using FIJI, and the statistical analyses were performed using GraphPad Prism.

### FM1-43FX (*N*-(3-Triethylammoniumpropyl)-4-(4-(Dibutylamino) Styryl) Pyridinium Dibromide) dye uptake assay

As described previously (*63*), the inner ears were harvested from the young and old mice and transferred to DMEM supplemented with 10% FBS, and the utricle and cristae were microdissected efficiently. The tissues were immersed in 10 μM FM1-43 (Invitrogen, Cat: F35355) for 1 min 30 seconds, followed by washes with DMEM. The tissues were fixed with 4% PFA for 20 minutes, washed with DMEM, and mounted. The images were acquired using Zeiss 980 confocal microscope. Images were processed using FIJI, and optical sections were made using Orthoslicer in Imaris. Fluorescence intensities were quantified using FIJI, and statistical analysis was performed using GraphPad Prism.

### Senescence-associated β-galactosidase (SA-β-gal) staining and imaging

According to the user manual, senescence-associated β-gal (SA-β-gal) staining was performed using senescence-associated β-gal (Cell Signaling #9860) staining kit. The temporal bones were fixed in the 1X fixative solution provided in the kit for 30 minutes, as previously described (*70*). Then the cochlea and the vestibular system were microdissected. According to the manufacturer’s protocol, 1x staining solution, X-Gal, and β-galactosidase staining solutions were prepared. The cochlear and vestibular whole mounts were incubated in the staining solution overnight at 37°C. Samples were mounted and imaged using Olympus VS120 virtual slide scanner.

## Acknowledgment

We thank Dr. Marisa Zallocchi for generously sharing the reagents. This research partially utilized the Auditory and Vestibular Technology (AVT) core (grants GM103427, GM139762), Histology and Molecular core facility (grants RRID:SCR_025163), Department of Chemistry & Biochemistry (EPSCoR) at Creighton University, and DNA Sequencing Core Facility at the University of Nebraska (grant RR018788).

## Author Contributions

Conceptualization: S.K., D.H.

Methodology: S.K., H.L., D.H.

Investigation: S.K., H.L., D.H., S.V., S.T., C.B., M.Z., B.J.B.

Visualization: S.K. Supervision: D.H., L.T., B.K.

Writing—original draft: S.K.

Writing—review & editing: S.K., D.H., B.K., L.T., C.B., S.V., B.J.B.

## Data and materials availability

Publicly available standard packages and algorithms were used for the analysis of our sequencing data and no custom code or algorithms were generated. All relevant data are included in this paper and the supplementary materials. The single-cell RNA sequencing data generated in this study have been deposited in the NCBI Gene Expression Omnibus (GEO) under the accession numbers GSE283534 (young CBA/J) and GSE283708 (old CBA/J).

