## Supplementary Figures 1-8 and Table 1 for "Hair Bundle Degeneration is a Key Contributor to Age-Related Vestibular Dysfunction"

Samadhi Kulasooriya et al.

Corresponding authors:
David Z. He
Samadhi Kulasooriya

**This PDF file includes:**

Figs. S1 to S8
Supplementary figure legends
Table S1

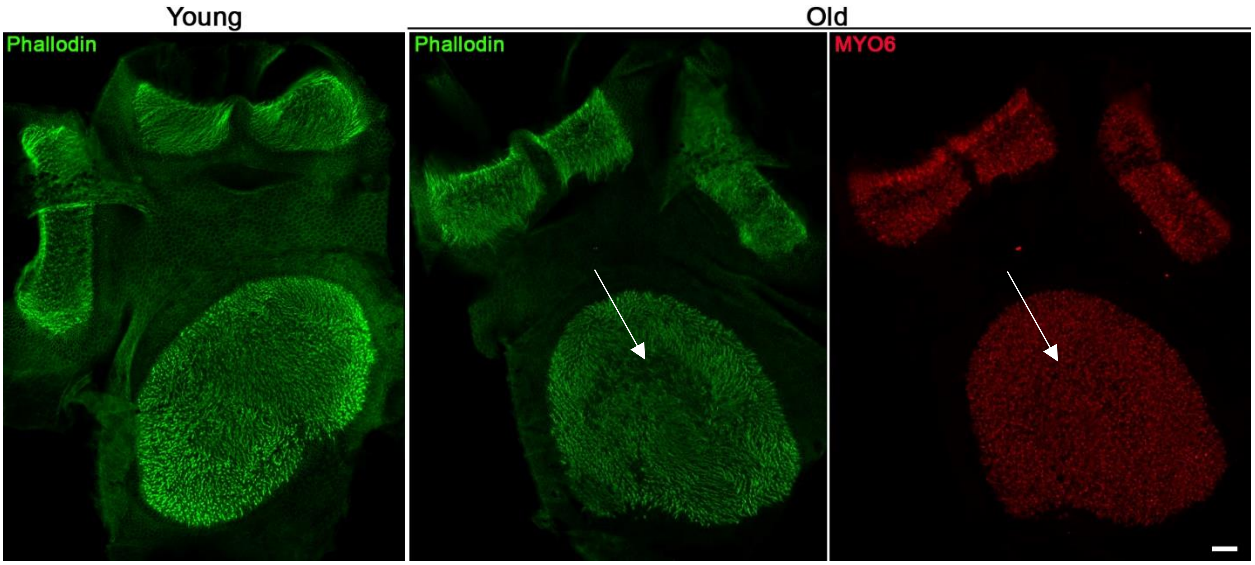


**fig. S1. Low magnification images from the young and old utricle and cristae.** Arrow indicates areas with bundle degeneration without loss of HCs. Scale bar, 50 μm.

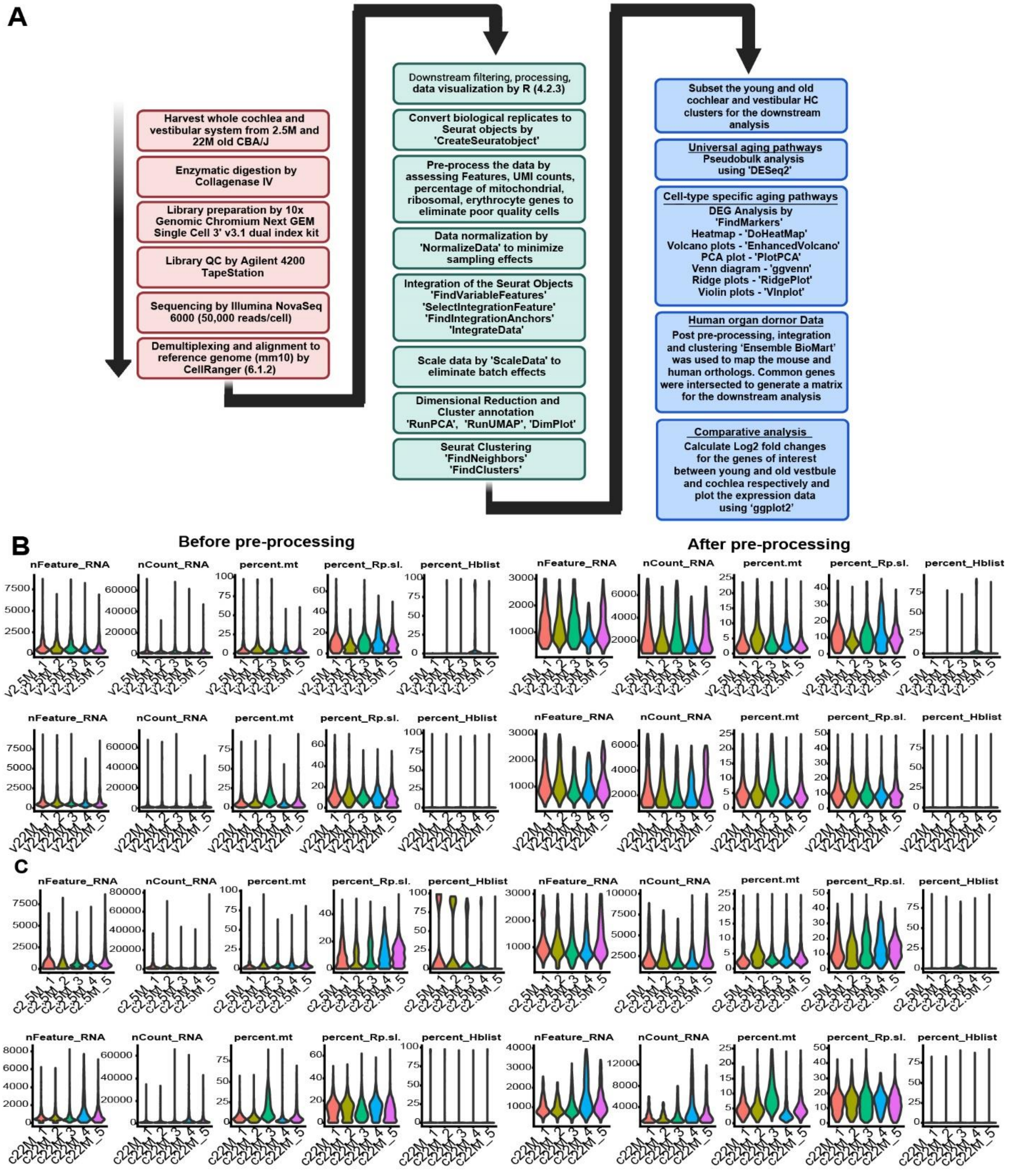


**fig. S2. Detailed scRNA-analysis workflow and pre-processing for quality control. (A)** The detailed computational workflow followed for the transcriptomic analysis. 2.5 and 22-month-old cochleae and vestibule were digested using collagenase IV, and sequencing libraries were prepared by 10x chromium next GEM single cell 3' v3.1, followed by QC assessment by Agilent 4200 TapeStation and sequencing by Illumina NoveSeq 6000 covering a depth of 50,000 reads/cell. Reads were demultiplexed and aligned by CellRanger. Downstream quality control and batch effect elimination were performed in R using Seurat and other packages. Once the clusters were annotated based on the known marker gene expression, HC clusters were subsetted from young and old samples to assess universal and cell-type specific aging signatures. Universal aging programs were evaluated via pseudobulk analysis, whereas cell-type specific aging programs were examined through DEG and GO/KEGG enrichment analysis. **(B and C)** Violin plots depicting filtering parameters such as features, counts, mitochondrial, ribosomal, and erythrocyte gene percentages. B indicates data before filtering and C indicates after filtering in young (v2.5M), and old (v22M) vestibule, and young (c2.5M) and old (c22M) cochleae respectively.

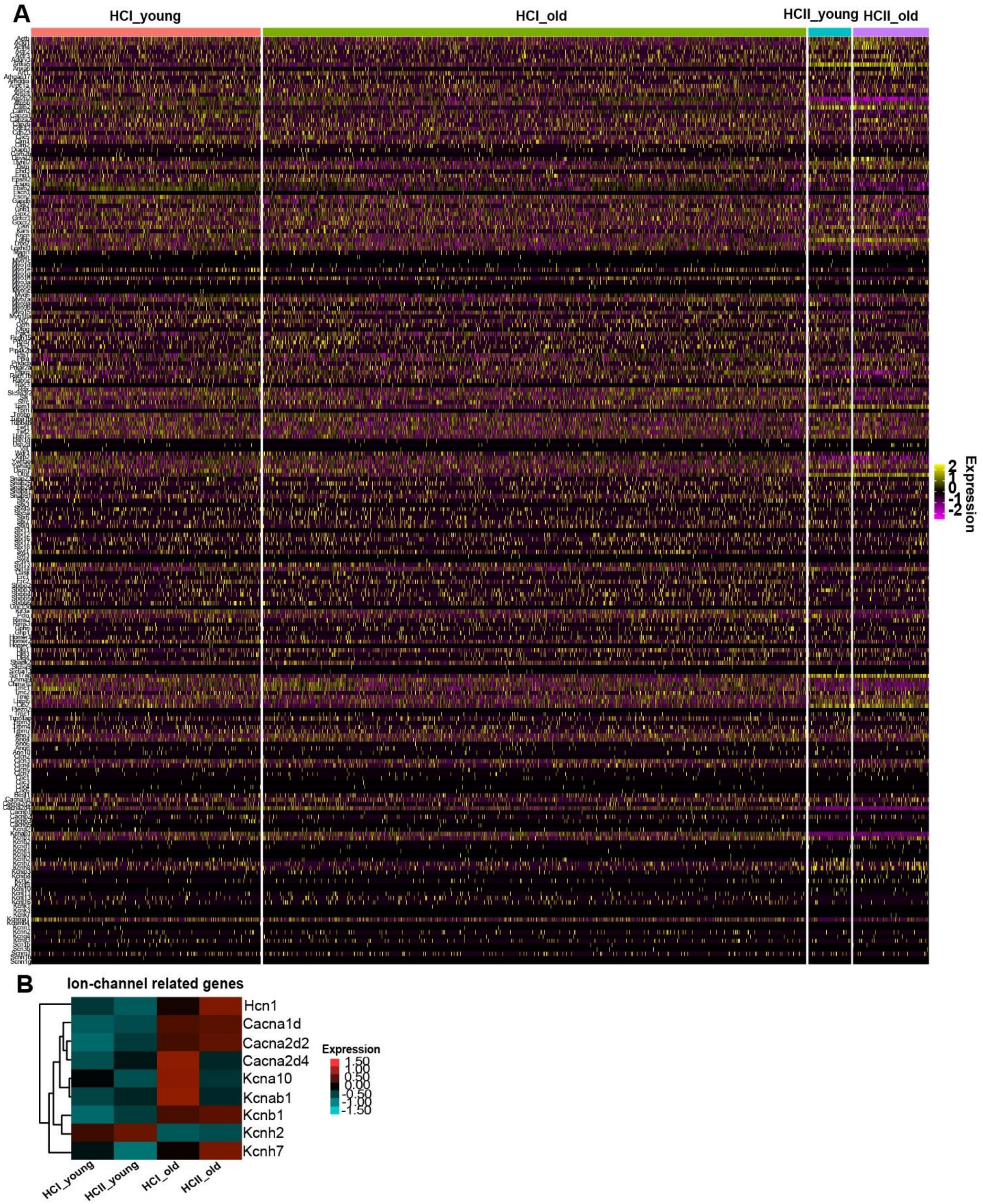


**fig. S3. (A)** HC-related gene (stereocilia, synapse and ion channel) expression in type I and II HCs at the individual cell level indicating heterogeneity. (**B)** Age-related changes in the key ion-channel related genes in vestibular HCs.

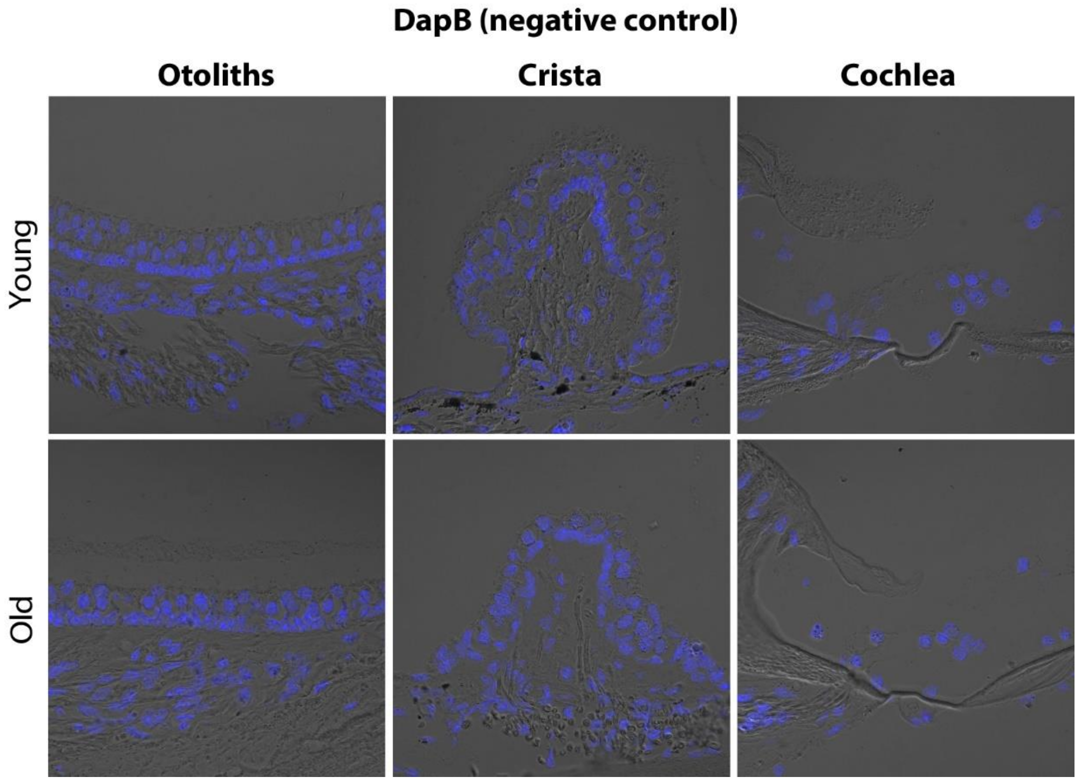


**fig. S4. (A)** Representative images of DapB (negative control) for RNAscope *in situ* hybridization. **(B)** Representative images of FM1-43 controls.

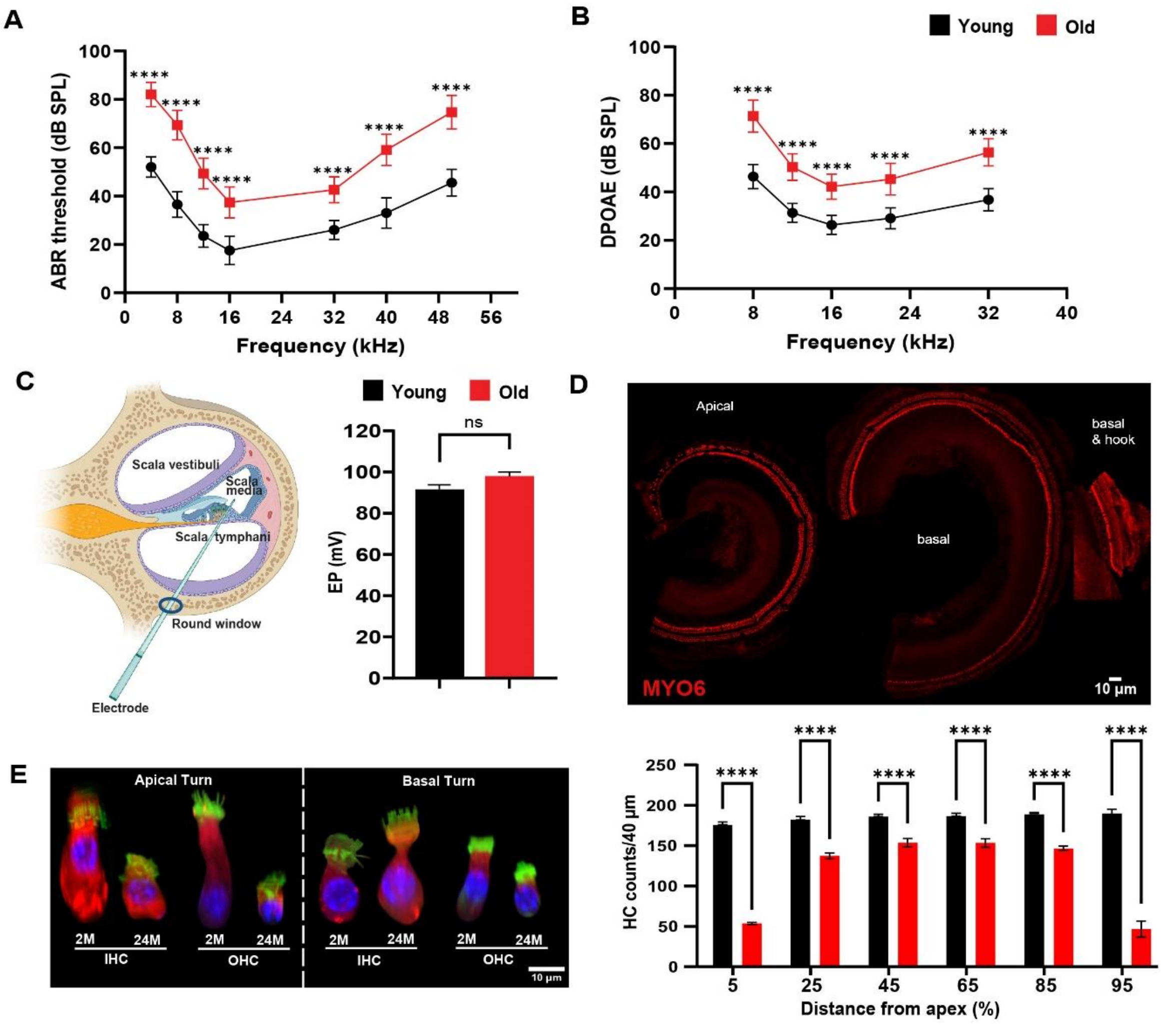


**fig. S5. Age-related auditory functional and morphological changes.** (**A**) Auditory brainstem response (ABR), **(B)** distortion product otoacoustic emissions (DPOAE) in young (n = 9) and old (n = 17) mice, and **(C)** endocochlear potential (EP) in young (n = 6) and old (n = 6) CBA/J. ABR and DPOAE were statistically analyzed by two-way ANOVA multiple comparisons. Data are shown as means SEM, (****p<0.0001, ns - non-significant. **(D)** Representative images of the cochlea immunolabeled with anti-MYO6 and the HC quantification along the tonotopic organization. **(E)** Confocal virtual sectionning images of individual HC morphology demonstrating cellular hypertrophy and atrophy in IHC and OHCs from apical and basal turns with aging. Virtual sections were acquired using the orthoslicer in Imaris, and the background was removed using Adobe Photoshop.

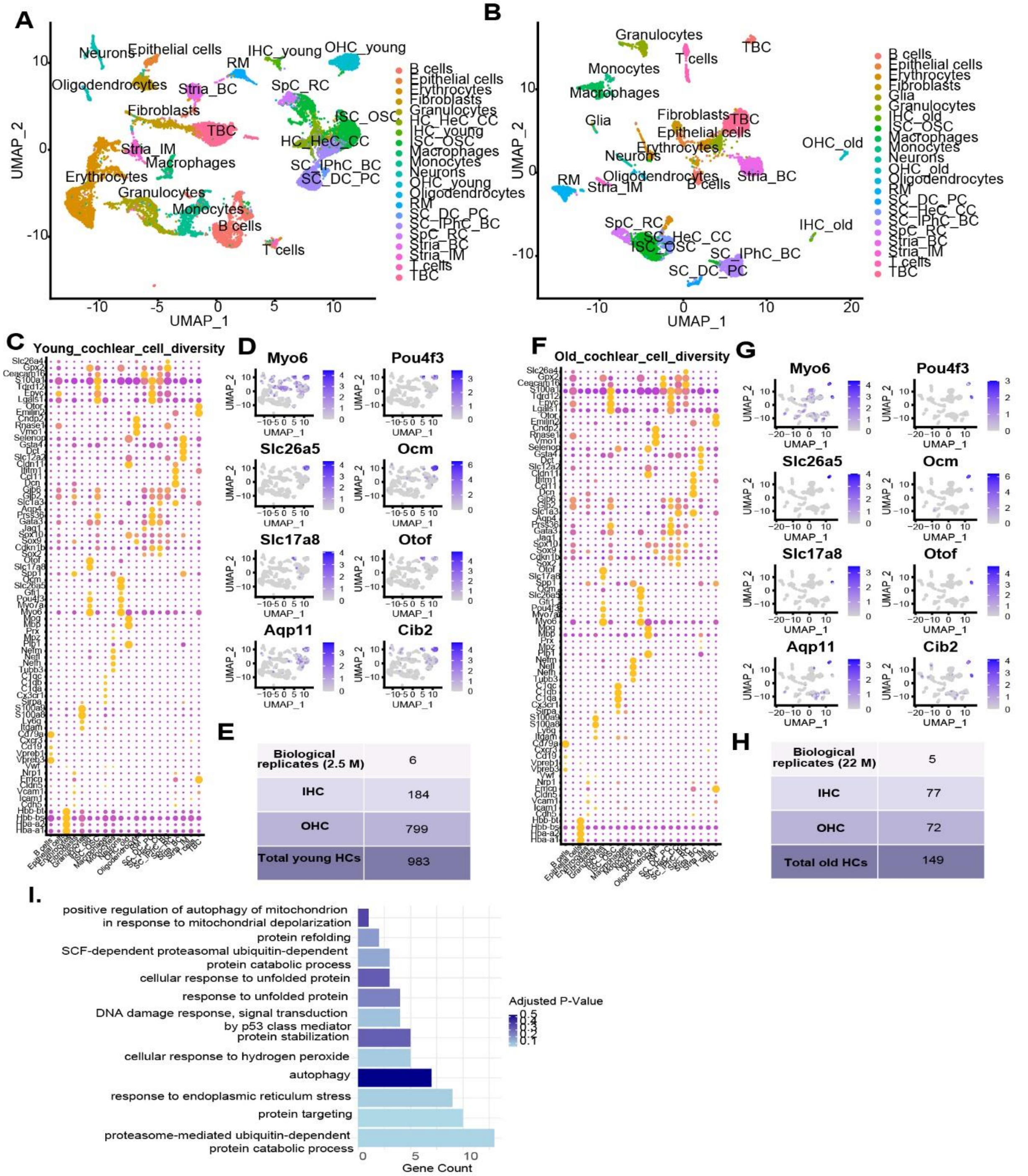


**fig. S6.** Young (2.5M) and old (22M) cochlear cell type distribution. (**A and B)** Uniform manifold approximation and projection (UMAP) plot shows the distribution of different cell types in young and old cochlear samples. (**C and F)** Dot plots indicating the expression of known marker genes used to identify the cell types and subsequent cluster annotation shown in B and C. The dot size represents the percentage of cells expressing the genes from each cluster, whereas the color indicates the expression level. Expression levels are normalized via z-score normalization. Thus, the average expression is zero, and positive or negative values indicate expression above or below average. (**D and G)** Feature plots demonstrating the HC-specific pan marker gene expression in the HC clusters identified in the UMAPs. (**E and H)** Number of inner and outer HCs identified from young and old cochlear samples for the downstream analysis. 7-8 mice were used per biological replicate, and a total of 5 biological replicates were used for the transcriptomic analysis. **(I)** Downregulated GO terms related to universal processes in old cochlear HCs compared to old vestibular HCS.

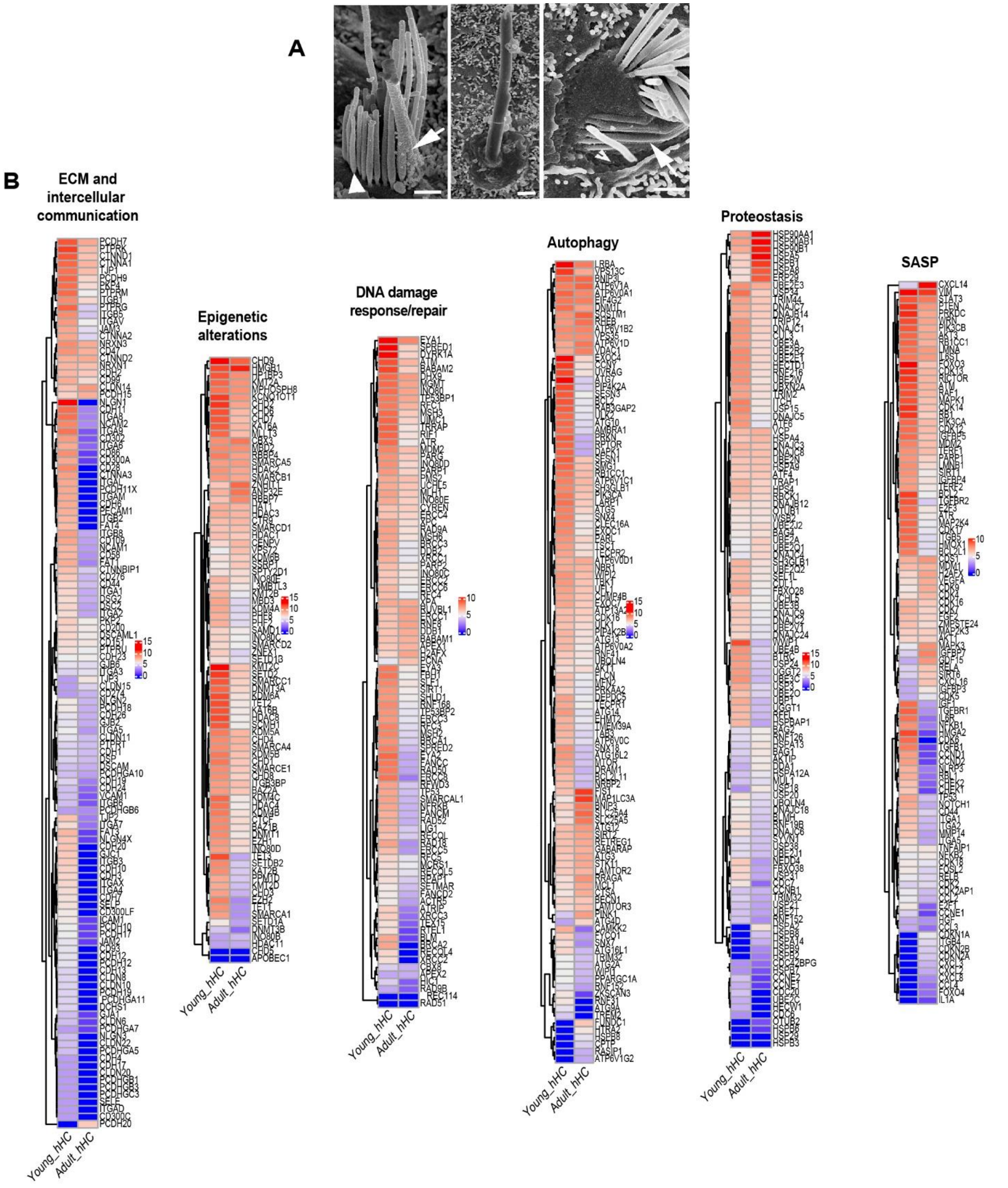


**fig. S7. (A)** Aging signatures in human vestibular HCs. **A.** Scanning electron micrographs indicating age-related stereociliopathy in humans (Adapted from Taylor et al, 2015, see ref. 71). Scale bar, 1 µm. **B.** Log2-transformed complex heatmaps of universal aging signatures in the human vestibular HCs. The color scale represents the pseudobulk Log2-normalized expression values. Red to blue indicates high to low expression.

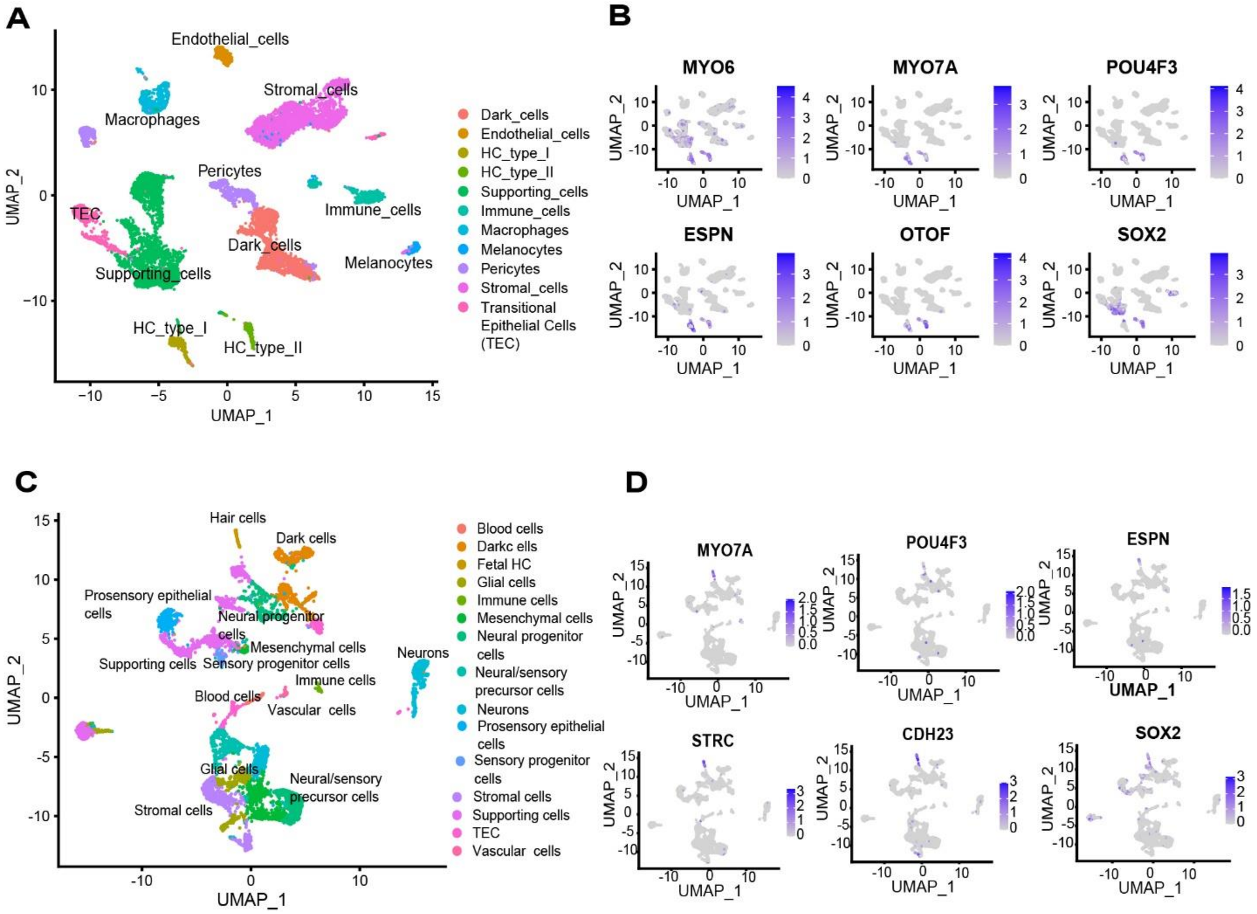


**fig. S8. (A)** Cell-type distribution of human organ donor transcriptomes**. A.** UMAP plot of fetal human vestibular tissue. **B.** UMAP plot of adult human vestibular tissues.

**Table S1: Antibodies and RNAscope probes**

| Antibody | Manufacturer | Catalog Number | Dilution |
|---|---|---|---|
| **Primary:** | | | |
| Anti-ESPN | Protein tech | 20717-1-AP | 1:200 |
| Anti-CCDC39 | Sigma Prestige Antibodies | HPA035564 | 1:200 |
| Anti- CCDC40 | Bioss | bs-8091R | 1:200 |
| Anti- MYO6 | Proteus | 25-6791 | 1:300 |
| Anti- MYO7A | Proteus | 25-6790 | 1:300 |
| Anti-acetylated-β-Tubulin | Sigma Millipore | T6793 | 1:200 |
| Anti-SOX2 | Invitrogen | 14-9811-82 | 1:300 |
| **Secondary:** | | | |
| Phalloidin FlourTM 488 | Invitrogen | A12379 | 1:200 |
| Phalloidin FlourTM Plus 405 | Invitrogen | A30104 | 1:200 |
| Goat anti-rabbit Fluor 568 | Invitrogen | A11011 | 1:200 |
| Donkey anti-mouse Fluor 568 | Invitrogen | A31570 | 1:200 |
| Goat anti-rabbit Fluor 488 | Invitrogen | A32731 | 1:200 |
| Goat anti-mouse Fluor 488 | Invitrogen | A11001 | 1:200 |
| **RNAscope Probes** | **Manufacturer** | **Catalog Number** | **Dilution** |
| *Espn* | Advanced Cell Diagnostics (ACD) Biotechne | 1061141 | N/A |
| *Tmc1* | | 520911 | |
| *Atp2b2* | | 1262061 | |
| *Fbxo2* | | 524151 | |
| *Pou4f3* | | 1740841 | |
| *Homer2* | | 581231 | |
| *Gjb2* | | 51881 | |
| *Hspb1* | | 488361 | |
| *Dapb* | | 310043 | |
